# Sequence-based prediction of condensate composition reveals that specificity can emerge from multivalent interactions among disordered regions

**DOI:** 10.1101/2025.06.13.659429

**Authors:** Jonas Wessén, Nancy De La Cruz, Heankel Lyons, Hue Sun Chan, Benjamin R. Sabari

## Abstract

While specificity of biomolecular interactions is typically understood to require interactions involving ordered structures, several biomolecular condensates exhibit specificity in the absence of apparent structural order. We have previously shown that condensates composed of the disordered region of MED1 partition specific proteins, mediated by sequence patterns of charged amino acids on the disordered regions of both MED1 and the interacting partner. Whether this specificity is due to an unknown ordered-structure-mediated interaction or from the dynamic multivalent interactions between the patterned charged amino acids in the disordered regions was unresolved. Here we show that a polymer physics-based model that only accounts for multivalent interactions among polymers in a statistical manner can largely explain published data on selective partitioning and make predictions that are subsequently experimentally validated. These results suggest that the specificity of condensate composition is underpinned to a substantial extent by multivalent interactions in the context of conformational disorder.

## Introduction

Specificity of biomolecular interactions is often conflated with folded or otherwise ordered structures. It has long been the goal of molecular biology to catalog the various ordered structures that biomolecules adopt and characterize how their interfaces engage with one another. The language and criteria used to define specificity of biomolecular interactions is informed by and assumes ordered-structure-mediated interactions or “site-specific interactions”, yet there is growing evidence that dynamic multivalent interactions involving intrinsically disordered regions (IDRs) can also exhibit specificity with functional outcomes as soluble complexes^1-9^ and in the context of meso-scale assemblies referred to as biomolecular condensates^10-14^. Theoretical examples of such a statistical mechanism of “fuzzy” molecular recognition afforded by IDRs include the sequence-dependent co-mixing and de-mixing of condensed polyampholyte species^13,15,16^ and other model IDRs^14,17^.

We recently demonstrated experimentally that condensates composed of a large IDR of MED1 (MED1^IDR^), the largest subunit of the transcriptional coactivator complex Mediator, selectively partitioned RNA Polymerase II (RNA Pol II) together with positive regulators of Pol II (e.g. SPT6) while excluding negative regulators of Pol II (e.g. NELFE)^10^. This functional specificity of MED1^IDR^ condensates required the sequence patterning of charged amino acids on disordered regions of partitioned partners into high local densities of either positive or negative charge, referred to as charge blocks. For example, alternating positive and negative charge blocks were found on the disordered region of SPT6 (SPT6^IDR^) but were absent from the disordered region of NELFE (NELFE^IDR^)^10^. Experimental manipulation of SPT6^IDR^ to remove charge blocks disrupted partitioning and manipulation of NELFE^IDR^ to add charge blocks promoted partitioning with functional consequences for RNA Pol II transcription. While these experiments provided evidence for the importance of sequence charge patterning in selective partitioning, it remained unresolved whether the observed specificity required an unknown ordered-structure-mediated interaction or were largely accounted for by dynamic multivalent contacts among the IDRs. Given the connection between this experimental work and theoretical treatments of sequence-dependent co-mixing and de-mixing of condensed polyampholytes^13,15,16^, here we test the extent to which a polymer theory-based statistical model for disordered chain molecules can explain our experimental data and make new testable predictions of MED1^IDR^ condensate specificity.

In the context of experimental observations that charge-patterning is required for specificity of MED1^IDR^ condensates, several theoretical approaches have been developed in recent years to account for various effects of sequence charge pattern on IDR behaviors. These efforts include using simulation and theory to investigate effects of sequence charge pattern on the conformational dimensions and phase separation propensities of polyampholytes (chain sequences with approximately equal numbers of positively and negatively charged units), leading to the recognition that sequence with more blocky charge patterns tend to adopt more compact conformations as an isolated chain^18,19^, and they also have a higher propensity to phase separate. The two effects appear to be significantly correlated though this correlation is not without limitations^20^. Sequence charge pattern also affects chain dynamics within condensates in molecular dynamics simulation^21^. Of particular relevance to the present investigation of IDR partitioning is that under the general field-theoretic framework, both approximate analytical theory^22^ and field-theoretic simulations^23-25^ have been applied to study sequence-specific effects in the mixing of condensed IDR species^15,16^. On the basis of these advances, we now introduce a novel, computationally efficient, and high throughput method derived from analytical polymer theory for predicting the partitioning of charged IDR sequences into a biomolecular condensate scaffolded by another charged IDR.

As detailed below, this high throughput method can be used to scan libraries of protein sequences to make predictions about the extent to which a protein sequence will (or will not) be partitioned into MED1^IDR^ condensates. After demonstrating that our method is capable of validating previously published experimental data, we make predictions for sequences for which there is no experimental data. Predictions of partitioning are then tested in cell-based assays where we can assess the degree of partitioning of any arbitrary sequence into a condensate of MED1^IDR^ tethered to a specific genomic locus. The cell-based assays show striking qualitative agreement with the model predictions and for the cases where the model disagrees with experimental results, provide quantitative insights into non-electrostatic contributions to selective partitioning in the MED1^IDR^ system, which may be incorporated into a more refined treatment in future studies. Taken together, these findings demonstrate, in the context of the experiments performed here, that selective partitioning can largely be explained by dynamic multivalent contacts among disordered regions without needing to invoke ordered-structure-mediated interactions.

## Results

### Rationale

Partitioning of IDRs into MED1^IDR^ condensates or any other IDR condensate is governed by multiple-chain interactions wherein the IDR and MED1^IDR^ chain conformations are expected to be disordered. Accordingly, pertinent biophysical properties that are thermodynamic in nature are determined by appropriate statistical averages over possible configurations entailing IDR-IDR, MED1^IDR^-MED1^IDR^, and IDR-MED1^IDR^ interactions. Based on the presumed predominance of these interactions in MED1^IDR^-scaffolded condensates, for simplicity our model does not account for possible interactions involving other molecular species that are present inside the cell. Here, as a first step, we are interested primarily in the effect of IDR sequence charge pattern on the electrostatics of partitioning, in which case the IDR-MED1^IDR^ interaction energy *U*_*ij*_ (where the label *i* stands for the IDR in question and the label *j* stands for MED1^IDR^ as in Eqs.1 and 2 below) is the usual screened-Coulomb potential given in Materials and Methods [Eq.6]. In view of the previously observed prominence of sequence charge pattern effects on IDR partitioning into MED1^IDR^ condensates^10^, it is conceptually worthwhile to first narrow our focus on probing how much electrostatics alone can physically rationalize such partitioning. Accordingly, while contributions from non-electrostatic interactions will clearly need to be considered in a more comprehensive approach, they will only be touched upon briefly in the present analysis.

Because the calculation of thermodynamic averages in principle entails considering all possible chain configurations, it is computationally challenging to study such a multiple-chain system using molecular dynamics simulations, especially if the goal is to assess the predicted IDR-sequence-dependent behaviors for many such systems for different IDRs. In lieu of taking into account all possible chain configurations, an intuitive understanding of the physics of sequence-dependent client-scaffold interactions can be attained by considering the constraining effects of chain connectivity on such interactions. As illustrated schematically in **Fig.1A**, among the many possible chain configurations in a client-scaffold condensate, chain connectivity makes it probable for amino acid residues that are sequentially local (i.e., close to one another along the chain sequence) on one intrinsically disordered protein chain to interact with sequentially local amino acid residues along one or more spatially neighboring chains. Accordingly, we propose an approximate, intuitive, semi-quantitative way to characterize the strength/weakness of partitioning-favoring client-scaffold interaction by computing a Boltzmann interaction energy averaged over different relative client-scaffold positions as the client sequence slides along the scaffold sequence aligned in both the parallel and antiparallel manners (**Fig.1B**). It is noteworthy that although this sequence-sliding analysis does not cover all possible chain configurations, it covers multiple scenarios of inter-chain interactions among sequentially local residues involving a pair or more than one pair of chains and/or non-interacting chain ends (examples 1—5 in **Fig.1B**).

**Fig. 1.**
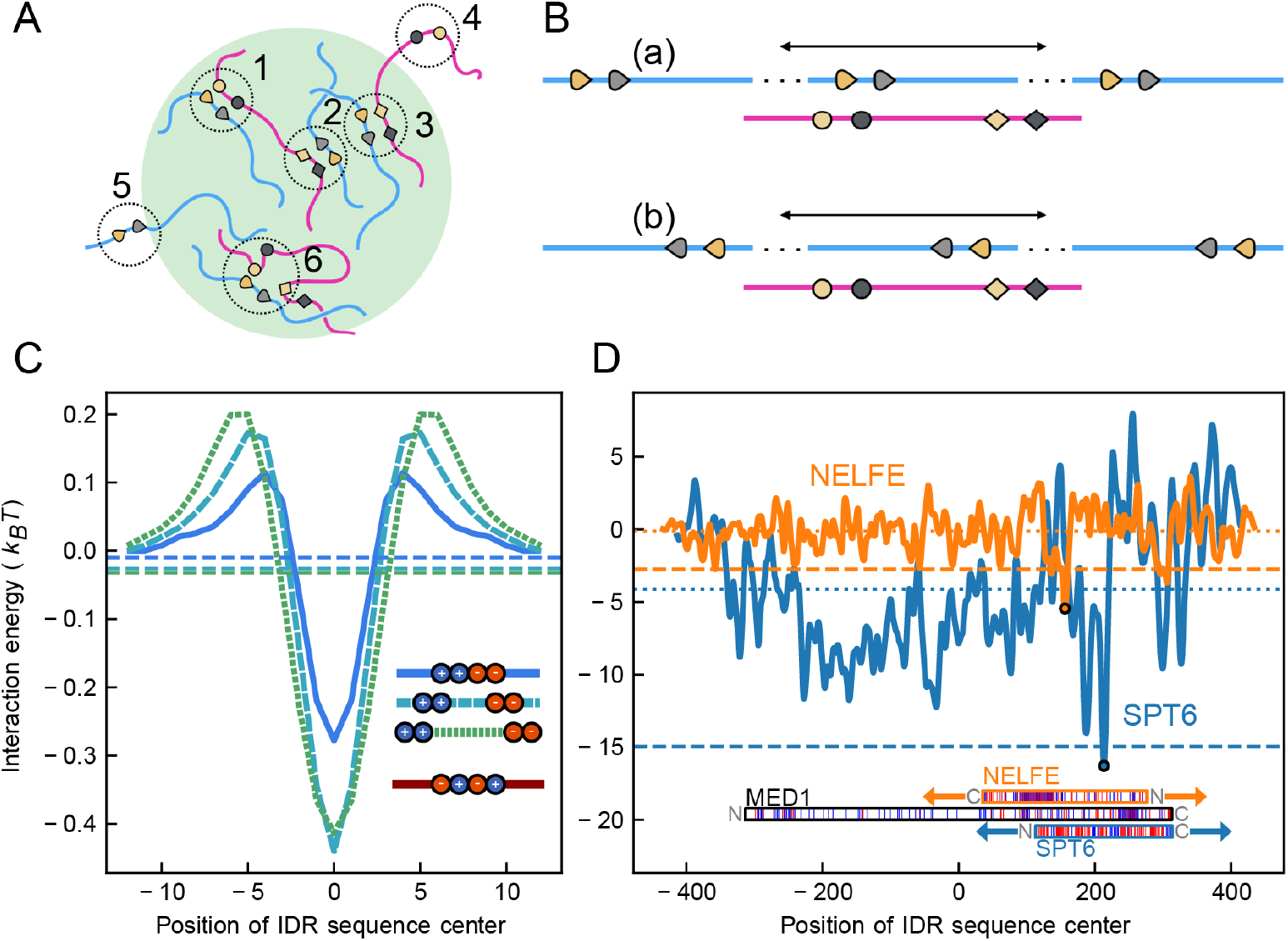
Physics of sequence-dependent IDR association and partitioning illustrated by an intuitive sliding-sequence consideration. **Fig.1A:** Schematics of interactions between an IDR scaffold (red chains) and an IDR client (blue chains) in and around a condensate (green area), wherein amino acid residues at selected positions along the two IDR sequences are represented by symbols of different shapes and colors. The dotted circles highlight several possible configurations of interchain interactions between the IDR sequences: (1) Because of chain connectivity, it is physically plausible for contact-like interactions to exist between chains that are locally approximately aligned; (2) in the condensed phase, these locally aligned contact-like interactions can be between different pair of chains at different sequence positions; and (3) the local alignments can be parallel or antiparallel with respect to the N- to C-terminus directions of the two IDRs. (4,5) More generally, it is also possible that some chain segments of either IDR may not be interacting in close spatial proximity with another IDR, especially when the segments are outside the condensed phase, and that (6) some contact-like interactions may not involve locally aligned chain segments at all. **Fig.1B:** The sliding-sequence analysis in the present work entails determination of interaction energies as the client sequence (blue) is moved along the scaffold sequence (red) as indicated by the double arrows. Three examples of different relative positions of the two sequences are schematically depicted in parallel as well as in antiparallel alignments (a,b).**Fig.1C:** Simple 8-bead model sequences each with four electric charges. Positive and negative charges are represented, respectively, by blue and red beads in the inset. The interaction of each of the three model client sequences (marked by blue, light blue, and green lines in the inset) with the model scaffold sequence (red line) is assessed by *U*_*ij*_/*k*_B_*T* (potential energy given by Eq.6 in units of *k*_B_*T*, vertical variable) as a function of the distance between the centers of the client and scaffold sequences (horizontal variable). The *U*_*ij*_/*k*_B_*T* energy profiles (curves) and the Boltzmann-weighted average energies ⟨*U*_*ij*_⟩/*k*_B_*T* (horizontal dashed lines) are shown using the same color code for the clients as that in the inset. **Fig.1D:** IDRs with a blockier charge pattern interact more favorably with the IDR of MED1. As in Fig.1C, the vertical variable is *U*_*ij*_/*k*_B_*T* and the horizontal variable is the distance between the centers of the MED1 and the NELFE or SPT6 IDR sequence. Positively and negatively charged residues are indicated by blue and red lines in the depiction of the MED1 and IDR sequences (bottom, with the N and C termini marked). The NELFE and SPT6 sequences are shown in their respective positions at which the interaction with MED1 is most favorable (most negative energies marked by circles along the orange and blue curves). Each of the dotted lines (same color code for the IDRs) is the arithmetic mean of the screened Coulomb energy (vertical variable) over the horizontal range; the thicker dashed curves are the Boltzmann-weighted average energies ⟨*U*_*ij*_⟩/*k*_B_*T*. Each of the energy profiles provided in Fig.1C and Fig.1D is for the alignment that has the lowest energy minimum. For presentational consistency, each of the averaged energies marked by the horizontal dashed and dotted lines is computed using the corresponding shown energy profile without contributions from the energy profile for the opposite alignment.

The sliding-sequence analysis is illustrated by the simple model sequences in **Fig.1C**. Each of these 8-bead model sequences have two positively charged, two negatively charged, and four electrically neutral residues. The top three sequences and the bottom sequence shown in the inset are taken, respectively, to be client sequences with different sequence separations between their positive and negative charge blocks and a scaffold with an alternating charge pattern in the middle of the sequence. For each model client sequence, the plotted curve in **Fig.1C** is the interaction energy *U*_*ij*_(*x*) between the client and scaffold sequences as a function of the horizontal variable *x* for the relative position of the two sequences in units of bond length *b*. A lower (more negative) *U*_*ij*_(*x*) value means a more favorable client-scaffold interaction at that position. Each of the horizontal dashed lines (same color code) is the Boltzmann-weighted energy ⟨*U*_*ij*_⟩/*k*_B_*T* ≡ Σ_*x*_ *U*_*ij*_(*x*) exp(−*U*_*ij*_(*x*)/*k*_B_*T*)/*k*_B_*T* Σ_*x*_ exp(−*U*_*ij*_(*x*)/*k*_B_*T*) averaged over the sliding range. *U*_*ij*_(*x*) is the position-dependent screened Coulomb energy in Eq.6 (Materials and Methods) applied to an inter-chain distance (perpendicular to the two chains) of 2.5*b* between the two sequences where *b* is the bond-length separation between successive beads along a chain which in turn corresponds to the C_α_-C_α_ virtual bond length (*b* ≈ 3.8 Å). In the above expression, *k*B is Boltzmann’s constant and *T* is absolute temperature. The choice of an inter-chain distance of 2—3*b* in **Fig.1C** and results of other sliding-sequence analyses presented below is motivated by our observation that this range of inter-chain distances is sufficiently small to allow for significant inter-chain interactions expected in a condensed phase but not too small to entail excessive fluctuations in *U*_*ij*_(*x*). Beside this consideration, the choice is to some extent arbitrary and, as such, should be regarded as tentative. It will be instructive to assess other inter-chain distances in future developments of similar sliding-sequence approaches. In any event, the simple examples in **Fig.1C** showcase the sequence-dependent nature of client-scaffold interactions. For the same given scaffold sequence, **Fig.1C** indicates that clients with a longer sequence separation between their charge blocks exhibit a more favorable (lower) Boltzmann-weighted average interaction energy (horizontal dashed lines) with the scaffold.

The above-described sliding-sequence analysis thus offers an intuitive physical picture as to how the charge sequences of an IDR and MED1^IDR^ may affect IDR partitioning by essentially using a set of “sliding-sequence’’ configurations as representative of the underlying, much more complex IDR-MED1^IDR^ interaction. This approach is next applied in **Fig.1D** to natural disordered regions from NELFE and SPT6 (sequences for all regions used in this study are provided in **Table S1**). These disordered regions are chosen as examples because we have previously determined experimentally that this disordered region of SPT6 has high partitioning while the disordered region of NELFE has low partitioning^10^. In the same vein as the simple examples of our sliding-sequence analysis in **Fig.1C, Fig.1D** illustrates the essential physics of charge-sequence-dependent IDR-MED1^IDR^ interactions by the position-dependent screened Coulomb energy (in units of *k*_B_*T*) *U*_*ij*_(*x*)/*k*_B_*T* between the disordered regions of MED1 and NELFE (orange lines). The same sliding-sequence analysis is applied to the N-terminal disordered region of SPT6 (blue lines) as each of the IDRs slides along MED1^IDR^ (indicated by the arrows), now at an inter-chain distance of 3 C_α_-C_α_ virtual bond lengths (3*b* ≈ 11.4 Å). Again, a lower (more negative) *U*_*ij*_(*x*)/*k*_B_*T* value means a more favorable IDR-MED1^IDR^ interaction. (Note that a slightly smaller *b* was used for the simple model sequences in Fig.1C to enhance differences in their inter-chain interactions for clarity because the model sequences are much shorter.) In addition to the Boltzmann-weighted thermodynamic averages introduced in **Fig.1C**, which are plotted in **Fig.1D** by thick horizontal dashed lines, horizontal dotted lines in **Fig.1D** provide also the arithmetic mean Σ_*x*_ *U*_*ij*_(*x*)/ *k*_B_*T* Σ_*x*_ 1 over the range of horizontal variable *x* considered. By comparison, although both the Boltzmann-weighted and arithmetic mean energies indicate consistently that SPT6 interacts more favorably with MED1 than NELFE, the Boltzmann average, which exhibits a larger difference in interaction energy between the disordered regions of SPT6 and NELFE, is expected to be more representative of the physical situation than the arithmetic mean because a basic tenet of statistical mechanics is that configurations that are energetically more favorable are populated more prominently in a constant-temperature ensemble and therefore should be given more weight in the thermodynamic average. Accordingly, the sliding-sequence analysis offers a physical rationalization for the experimental observation that SPT6^IDR^ partitions significantly more strongly into MED1^IDR^ (i.e., has a higher partition coefficient, PC) than NELFE^IDR 10^ because the results in **Fig.1D** shows clearly that SPT6 interacts significantly more favorably with MED1 than NELFE when averaged over the sliding-sequence configurations.

Nonetheless, while the sliding-sequence analysis offers semi-quantitative predictions for PC (a potentially useful extension to include non-electrostatic interactions will be discussed below), the procedure is time consuming in general because for a pair of sequences of length ∼*N*, it requires ∼*N*^3^ energy calculations. Molecular dynamics simulation of the partitioning process is even more computationally expensive. Another possible theoretical avenue is to apply random phase approximation (RPA) formalism^24,26,27^ to analytically model the binary phase separation of an isolated system containing an IDR species and MED1^IDR^ on equal footing. An exploratory calculation for wildtype (WT) and two variants of SPT6^IDR^, which were experimentally determined to have reduced partitioning^10^, produced a trend of predicted PCs consistent with the experimentally observed trend, supporting RPA as an effective means to capture the effect of sequence charge pattern on IDP interactions (SI text and **Fig.S1**). However, as it stands, the application of this method to PC prediction is limited because the model assumptions are too restrictive for quantitative match with experiments, and it is numerically costly for any task that requires a high throughput.

### The FH-RPA model

As described in Materials and Methods and detailed in SI, we found that experimental PCs can be rationalized and predicted quantitatively to a substantial degree through capturing the physics of IDR-MED1^IDR^ interaction approximately by treating MED1^IDR^ as the scaffold of a preformed condensate and the IDR as a client species. In this approach, the theory-predicted PC for IDR type *i* into MED1^IDR^ (labeled as *j*) is governed by a Flory interaction parameter^28^ *χ*_*ij*_:

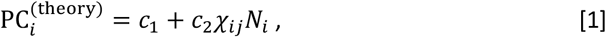

where *N*_*i*_ is the chain length (number of amino acid residues) of the IDR client *i, c*_1_ and *c*_2_ are empirical coefficients to be determined by fitting this formula to experimentally measured PCs. When only electrostatic interactions are taken into account, *χ*_*ij*_ can be approximated as a sum of a zeroth order term 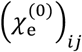 that depends only on the net e *ij* charges of the IDR and MED1^IDR^, and a first order random phase approximation (RPA) term

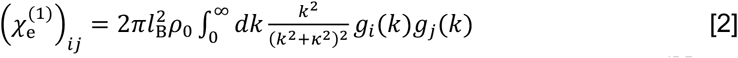

that takes into account the sequence charge patterns of the IDR and MED1^IDR^. In Eq.2, *l*_B_ is the Bjerrum length that determines the strength of electrostatic interactions in a thermal environment, *ρ* _0_ is a reference density for the solution system, *κ* is the inverse Debye screening length, and *g*_*i*_(*k*), a function of reciprocal space (Fourier-transformed) coordinate *k*, is dependent on the sequence charge pattern and involves a smearing length *a* for the IDR residues in the field-theoretic description. As will be discussed in more detail below, charge patterns involving residue positions far apart along the chain sequence (i.e., nonlocal patterns encompassing charges separated by long contour lengths) are only captured by *g*_*i*_(*k*) with small *k* values, whereas charge patterns involving residue positions close (local) to one another along the chain sequence are characterized by *g*_*i*_(*k*) with both small and large *k* values (expressions for *l*_B_ and *g*_*i*_(*k*) are provided in Materials and Methods). Because this formalism combines Flory-Huggins (FH) mean-field theory (MFT)^28^ with RPA treatment of sequence-pattern-dependent electrostatic interactions of charged polymers^24,26,27^, we refer to it as the FH-RPA method (Eqs.1, 2, 4-9).

It follows from Eq.2 above that the sequence-dependent RPA contribution to the Flory-Huggins interaction parameter *χ*_*ij*_ in Eq.1 satisfies the proportionality relation

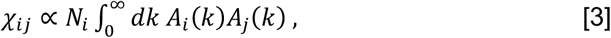

where

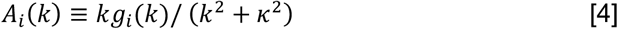

and *g*_*i*_(*k*) is proportional to a sum of terms in the form of

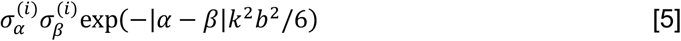

in which 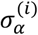 is the charge at sequence position *α* for sequence types *i* (Eq.6 in Materials and Methods). The FH-RPA approach is illustrated in **Fig.2A** using the simple model scaffold sequence (red curve) and the three model client sequences (blue and green curves) introduced in Fig.1C. A clear trend is seen in **Fig.2A** in that the *A*_*i*_(*k*) profile of a client sequence increases (i.e., takes higher values) with increasing sequence separation between the charge blocks (i.e., when the sequence charge pattern is more “blocky”).

**Fig. 2.**
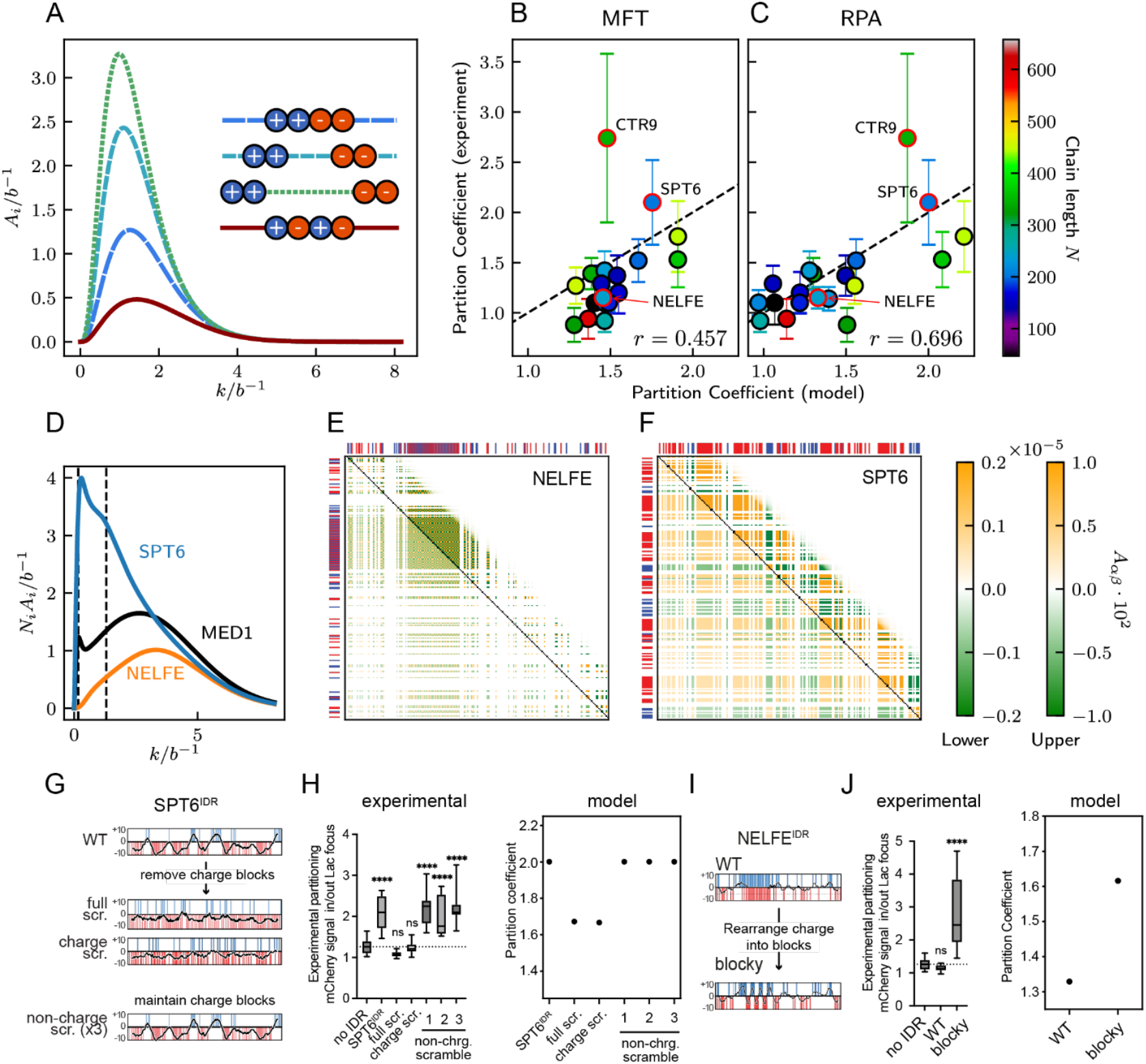
FH-RPA model rationalizes effects of charge scrambling and other sequence charge pattern variations on IDR PCs. **Fig.2A:** The FH-RPA account of sequence-dependent client-scaffold by *N*_*i*_*A*_*i*_(*k*) [Eq.1 and Eq.2] is illustrated by the *A*_*i*_(*k*) curves of the three simple model client sequences and the model scaffold sequence (inset) introduced in Fig.1C (same color code). Note that the *N*_*i*_ factor is omitted here in the vertical variable because all four sequences considered are of the same length. **Fig.2B,C:** Scatter plots of published experimental versus theory-predicted PCs for the partitioning of 19 natural (wildtype, WT) client sequences into MED1 (excluding MED1 into MED1) used in our parameter optimization. Correlation is significantly enhanced in RPA (C) relative to MFT (B); *r* is Pearson correlation coefficient. Sequence chain lengths are color coded. Experimental values (symbols) are presented as the average of 10 measurements. The corresponding standard deviations are indicated by the experimental error bars (values are provided in Table S2). **Fig.2D:** The FH-RPA-predicted sequence-dependent PC of an IDR into MED1 condensates is governed by *N*_*i*_*A*_*i*_(*k*) [Eq.1 and Eq.2]. **Fig.2E,F:** The NELFE (E) and SPT6 (F) *A*_*αβ*_(*k*), *α* ≠ *β* heat maps for the smaller and larger *k* values [dashed lines in (D)] are provided, respectively, by the lower and upper triangles. White areas represent *A*_*αβ*_(*k*) = 0 or *A*_*αβ*_(*k*) ≈ 0 (right scales). Sequence charge patterns of NELFE and SPT6 [as in Fig.1D] are shown along the heat map axes. **Fig.2G:** Representation of the charge patterning and net charge per residue (NCPR) profile of WT SPT6^IDR^ and the indicated sequence variants. The position of acidic or basic amino acids is indicated as vertical red or blue lines, respectively. The horizontal black curve represents the NCPR profile using a 10 amino acid sliding window, wherein the net charge of successive 10-residue windows is plotted (10 times the net charge per residue). Consequently, each NCPR profile has a possible max of +10 and a possible min of -10 centered at 0 charge. **Fig.2H:** Left: compilation of previously published experimental data of the partitioning of fluorescence signal from mCherry fused to the indicated control or SPT6^IDR^ variant inside and outside of CFP-LacI-MED1^IDR^ foci in Lac array cells. Data are presented as a boxplot (min-max). p-value, one-way ANOVA with multiple comparison test vs. the no IDR control. Ordinary one-way ANOVA with Dunnet’s multiple comparisons test to a “no IDR” mCherry control (N, 11) was performed for WT SPT6^IDR^ (N, 10; p, <0.0001), full scramble (N, 10; p, 0.6742), charge scramble (N, 10; p, >0.9999), and three independent non-charge scrambles (each with N, 10; p, <0.0001). These data were originally reported by^10^. Dashed horizontal line indicates median of the “no IDR” control. Right: FH-RPA model predictions of partition coefficient values (Eq.1) for indicated SPT6^IDR^ variant into MED1^IDR^ condensates. **Fig.2I:** Similar to 2C, but for NELFE or a charge rearranged variant designed to mimic the blocky charge patterning found in SPT6^IDR^.**Fig.2JF:** Similar to 2D, but for NELFE WT and charge rearranged variant (blocky). Ordinary one-way ANOVA with Dunnet’s multiple comparisons test to a “no IDR” mCherry control (N, 11) was performed for WT NELFE^IDR^ (N,10; p, 0.8805) and blocky (N, 10; p, <0.0001).

According to Eq.3 above, because the model client sequences share the same chain length *N*_*i*_ (= 8), a sequence with a higher *A*_*i*_(*k*) profile is predicted by RPA to have a higher PC, and this trend is consistent with the trend in Fig.1C from sliding-sequence analysis.

While recognizing that the exact mathematical forms in Eqs.3-5 follow rigorously from the RPA derivation in Materials and Methods and SI, it is also straightforward to see from the expression in Eq.5 why more blocky sequence patterns result in larger *g*_*i*_(*k*) and thus larger *A*_*i*_(*k*) as exemplified in **Fig.2A**. Because of the 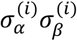 factor in Eq.5, a pair of charges of the same sign at the *α*th and *β*th positions contributes positively to *A*_*i*_(*k*) whereas a pair of charges of opposite signs contributes negatively. When the sequence charge pattern is blocky, for 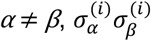 tends to be positive for small |*α* − *β*| because sequence-local charges tend to be of the same sign, and 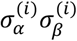 tends to negative mostly for large |*α* − *β*| when *α* and *β* are sequence positions in oppositely charged blocks (note that α = β terms are always positive). Since the value of the exponential factor in Eq.5 (which is always positive) decreases with increasing |*α* − *β*| for *k* > 0, it follows that blocky charged sequences have larger positive contributions compared to the magnitudes of negative contributions and thus resulting in larger *A*_*i*_(*k*)s. Conversely, for sequences with a well-mixed charge pattern, 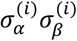 tends to change sign for small variations in |*α* − *β*| and therefore terms in the form of Eq.5 tend to cancel partially, resulting in smaller *A*_*i*_(*k*)s.

In light of this consideration, the FH-RPA prediction that PC increases linearly with *χ*_*ij*_*N*_*i*_ (Eq.1) which in turn contains an RPA term proportional to the integral of the product in Eq.3 of *A*_*i*_(*k*) for the client and *A*_*j*_(*k*) for the scaffold may be understood, at least partially, as a statement that partitioning is favored by a blocky client as well as a blocky scaffold. For client and scaffold sequences with substantial numbers of both positively and negatively charged residues such that they can broadly be viewed as polyampholytes for which RPA is best suited^29^, this physical picture makes intuitive sense because a blocky client and a blocky scaffold would allow for pairing of oppositely charged blocks on the client and the scaffold that are strongly favorable and thus resulting in higher PCs, whereas a less blocky client or a less blocky scaffold or both would lead to weaker interactions and thus smaller PCs. In the FH-RPA formulation, effects of any net positive-negative charge imbalance on the client and the scaffold are then accounted for by the zeroth order mean-field term that depends only on the net charges of the two sequences (Eq.8 in Materials and Methods). Beyond this general charge “blockiness” consideration, sequence-dependent client-scaffold interactions are also approximately taken into account by coupling their length scale-modulated sequence charge patterns at the same length scale (i.e., multiplying *A*_*i*_(*k*) and *A*_*j*_(*k*) at the same *k*) and integrating over all length scales (integrating over *k* in Eq.3). While this formulation does not account for all manners of interactions between the client and scaffold (in this sense it is similar to the sliding-sequence analysis), it is the essence of RPA which has proven to be useful for physical modeling of sequence-dependent phase separation^22^.

As an initial validation test and parameter optimization of FH-RPA, we used the experimentally measured PCs of 31 IDR sequences into MED1^IDR^ condensates (**Table S2** and **Fig.S2**) to maximize their Pearson correlation coefficient *r* with the term *χ*_*ij*_*N*_*i*_ in the theoretical Eq.1 by optimizing *l*_B_, *κ*, and *a* as fitting parameters (SI text and **Fig.S3**). The numerical values of *c*1 and *c*2 (> 0), which do not affect *r*, were subsequently determined by a linear fit to Eq.1 (SI text). This set of 31 partitioning measurements are compiled from our previous publication^10^ and include natural IDR sequences, variants of natural IDR sequences, and synthetic designed sequences. These partitioning measurements are all derived from the same cell-based fluorescence microscopy method using a U2OS cell line with large repeats of LacO DNA elements integrated into its genome^30^. Due to the high affinity interactions of LacI protein to LacO DNA, expression of a CFP-tagged LacI fused to MED1^IDR^ (CFP-LacI-MED1^IDR^) leads to a high local concentration of MED1^IDR^ at the LacO repeat locus creating a synthetic condensate which can be detected using confocal fluorescence microscopy as a bright CFP focus. To measure partitioning, we introduce a second protein sequence fused to mCherry and measure mCherry fluorescence signal inside and outside the CFP-LacI-MED1^IDR^ focus. This strategy has been used in many publications to assess partitioning in cells^10,12,31-36^.

This first exercise with the FH-RPA model indicates that sequence charge pattern plays a prominent role in determining IDR partitioning into MED1^IDR^ condensates. **Fig.S4** shows that if only the net charges of the IDRs are considered (MFT, using only the zeroth order term), the Pearson correlation coefficient is merely *r* ≈ 0.35 between theory-predicted and experimental PCs, whereas the correlation improves to *r* ≈ 0.67 when sequence charge patterns are taken into account (RPA, using both the zeroth and first order terms). These correlation coefficients are obtaining by practically treating *l*_B_, *κ*, and *a* as merely mathematical parameters to be optimized. To assess the robustness of our approach, we tested an example system with a set of experimentally accessible conditions with aqueous relative permittivity *ε*_r_ = 78.5, *T* ≈ 280 K (*l*_B_ = 2*b*) and Debye screening length *κ*^−1^ ≈ 3.8 nm (*κ* = 0.1*b*^−1^, i.e., ionic strength ∼ 6 mM), for which *r* ≈ 0.65 was obtained for FH-RPA (see SI), which is only slightly smaller than the *r* ≈ 0.67 obtained using optimized parameters, indicating that the correlation is not highly sensitive to parameter choice and thus putting the reasonably good correlation seen in our FH-RPA approach on a sound physical basis. We have further assessed the robustness of the obtained correlations by calculating the correlation coefficient *r* using the optimized parameters for every one of 500,000 samples of 19 randomly selected sequences (with replacements) out of the 19 wildtype (WT) client IDR sequences in **Table S2** (excluding MED1 itself). The resulting distributions in *r* for MFT and RPA, which are provided in Fig.S5, show a wide distribution for MFT (*Z*-score ≈ −0.280) but a significantly narrower distribution for RPA (*Z*-score ≈−0.252). Notably, virtually all *r* values in the RPA distribution are larger than 0.5, indicating that the reasonably good correlation between experimental and FH-RPA-predicted PCs is a robust observation.

Using the optimized parameters, the correlation between experimental and theory-predicted PCs for the subset of 19 natural (WT) client IDRs is shown in **Fig.2B,C**, again exhibiting improved correlation when sequence charge pattern is taken into account. Uncertainties in experimentally measured PCs are provided in **Fig.2B,C** by the error bars.

It is notable that outliers in the scatter plots tend to have larger experimental uncertainties, suggesting a tantalizing possibility that experiment-theory correlation—especially for RPA—may be further improved with more accurate measurements in the future. In any event, these results taken together demonstrate the utility of the FH-RPA model in predicting PC values for a relatively large library of IDRs. The results also demonstrate how combining mean field theory together with random phase approximation yields predictions that have greater agreement with experimental data.

### FH-RPA model rationalizes effects of charge scrambling on IDR PCs

To illustrate how FH-RPA captures sequence charge pattern-dependent IDR-MED1^IDR^ interactions, we display the quantity *N*_*i*_*A*_*i*_(*k*) ≡ *N*_*i*_*kg*_*i*_(*k*)/ (*k*^2^ + *κ*^2^) in **Fig.2D** for MED1^IDR^ and for the IDRs of NELFE and SPT6. These IDRs are chosen as examples of well partitioned (SPT6) and poorly partitioned (NELFE) sequences. Since 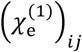 is the main factor in the FH-RPA-predicted PCs (Materials and Methods) and 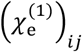 is proportional to 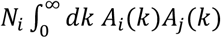 in accordance with the above Eq.2, the larger theory-predicted PC for SPT6 is seen here as a result of the larger *N*_*i*_*A*_*i*_(*k*) value of SPT6^IDR^ over that of NELFE^IDR^ for the entire range of *k* plotted in **Fig.2D**. How this difference arises from the two IDRs’ sequence charge patterns is further illustrated by the heat maps in **Fig.2E and F** for the quantity 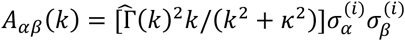 exp(−|*α* − *β*|*k*^2^*b*^2^/6) which is the smeared version of the quantity displayed in Eq.5 above [multiplied by the smearing factor 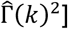, at one smaller and one larger *k* values. *A*_*αβ*_(*k*)s are individual terms contributing to 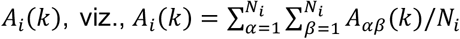. As discussed above, more blocky charged sequences have larger *A*_*i*_(*k*) values because of the positive *A*_*αβ*_(*k*) terms within a block of residues with mostly the same charge. The heat map for NELFE in **Fig.2E** exhibits mostly negative *A*_*αβ*_ values (green areas) and few positive *A*_*αβ*_ values (orange areas) because the NELFE sequence charge pattern (shown along the left and top sides of the heat map) is not blocky. In contrast, the heat map for SPT6 in **Fig.2F** exhibits mostly positive *A*_*αβ*_ values (orange areas) and fewer negative *A*_*αβ*_ values (green areas) because the SPT6 sequence charge pattern is blocky, as it can be seen clearly from **Fig.2F** that the positions of the orange areas in the heat map largely coincide with the positions of the sequence charge blocks along the left and top sides (negative charge blocks indicated by densely-spaced red lines). Accordingly, the larger *A*_*i*_(*k*) values of SPT6^IDR^ than those of NELFE^IDR^ arise from the significantly larger positive orange-colored area in the heat maps for SPT6^IDR^ than that for NELFE^IDR^. The heat maps in **Fig.2E** and **2F** also illustrate that *A*_*i*_(*k*) for small *k* characterizes both sequentially local (small |*α* − *β*|) and nonlocal (large |*α* − *β*|) charge patterns, as orange and green dots representing nonzero *A*_*αβ*_(*k*) values are present extensively in the lower triangles for small *k*. By comparison, orange and green dots are restricted to a narrow region near the diagonal of each of the upper triangles for large *k*, indicating that *A*_*αβ*_(*k*) ≈ 0 when |*α* − *β*| is larger than a relatively small value. Therefore, as mentioned above, only sequentially local charge patterns are characterized by *A*_*i*_(*k*) with large *k*.

As an illustration of how FH-RPA can discriminate between sequence patterning in qualitative agreement with previously published experimental data, we focused on sequence rearrangement experiments of SPT6^IDR^ or NELFE^IDR^. In these experiments, overall sequence composition, and therefore average charge properties, are maintained but the positions of residues are manipulated in either sequence to remove, maintain, or introduce charge patterning associated with partitioning into MED1^IDR^ condensates. For SPT6^IDR^ we made three different types of scrambled sequences (**Fig.2G**). Two of these remove blocky charge patterning by either scrambling all amino acids (full scramble) or scrambling only charged amino acids leaving non-charged amino acids unchanged (charge scramble). In the third type of sequence rearrangement, we scrambled the sequence of non-charged amino acids leaving charge amino acids unchanged thereby maintaining the wildtype charge patterning. This “non-charge scramble” was designed in three random iterations. Previously published^10^ results compiled here are in qualitative agreement with changes to PC derived from FH-RPA, both showing that removing blocky charge pattern disfavors partitioning and manipulations that maintain blocky charge pattern do not affect partitioning (**Fig.2H)**. For NELFE^IDR^, we rearranged charged amino acids to introduce charge blocks (**Fig.2I**) leading to an increase in partitioning measured experimentally and to an increase in PC values derived from FH-RPA (**Fig.2J**). Taken together, the FH-RPA model is able to explain changes in partitioning due to manipulations of sequence patterning even when the sequences have the same average charge properties. Notably, the MFT approach will fail in this regard because the MFT-predicted PC depends only on the amino acid composition of the sequence and therefore MFT always predicts the same PC for all scrambled versions of a given sequence.

### FH-RPA model successfully identifies high-PC regions in proteins

In view of the significant dependence of predicted PCs with IDR chain length (SI text) and the possibility of IDR chain-length-dependent reduction in conformational entropy upon partitioning into a condensed polymeric environment^37^, we next conducted a focused test of the FH-RPA model by considering IDR sequence segments sharing the same length. We do so by computing theory-predicted PCs for sliding sequence windows of 200 residues. The protocol, as described in **Fig.3A**, entails sliding a 200-residue window (grey bar, horizontal arrows) along a full-length sequence (human CTR9 in this example) and compute a theory-predicted PC at every step. This procedure produces a PC profile for the full-length sequence (black curve). To identify a high PC region within a full-length sequence, we first locate the position of the window with the maximum predicted PC (indicated by top horizontal red dashed line) and then take the two windows with 90% of the maximum predicted PC as the boundary of the high-PC region (horizontal orange bar in **Fig.3A**).

**Fig. 3.**
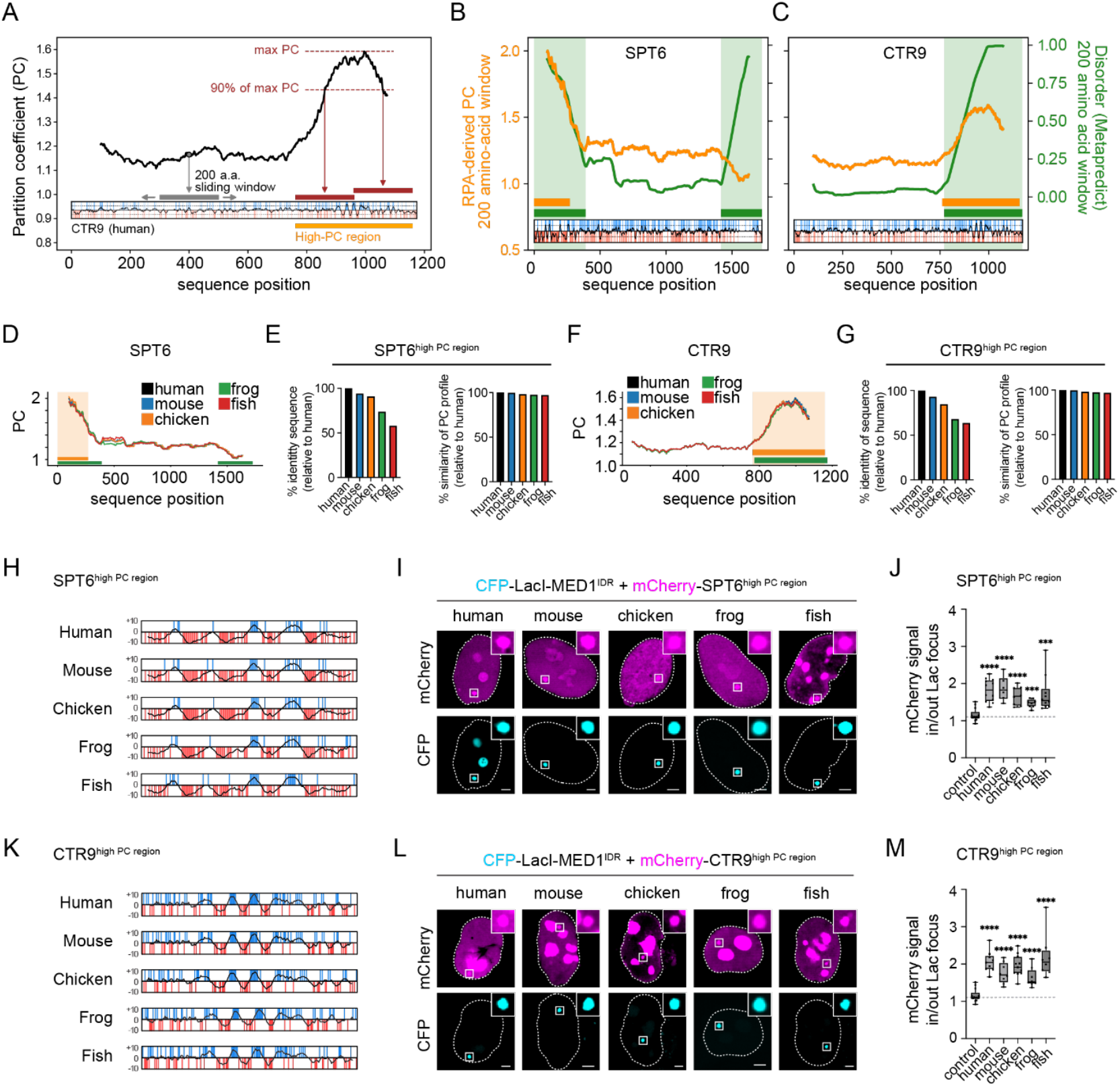
FH-RPA model successfully identifies high-PC sequence windows in CTR9 and SPT6. **Fig.3A:** Sliding-window analysis. Using human CTR9 as an example, PCs are computed for a 200-residue window (grey horizontal bar) sliding along (horizontal grey arrows) the full-length sequence and plotted at the midpoint of the sequence window (black curve in the main plot). The inset depicts the sequence charge pattern of the entire CTR9 as in Fig.1D, now with the black curve inside the inset showing sliding NCPR over consecutive 10-residue windows (inset vertical scale for charge ranges from −10 to +10 as in Fig.2G). The predicted high-PC region (orange bar) is constructed as described in the text; the red bars are the first and last 200-residue windows in the high-PC region. **Fig.3B:** Sliding-window analysis for human SPT6. PC profile is plotted as in Fig.3A, now in orange (left vertical scale), together with the Metapredict disorder profile computed for the same set of 200-residue windows (green curve, right vertical scale). The disorder regions, defined by disorder measure > 0.5, are shaded in green and indicated by the horizontal green bar. The high-PC regions are indicated by the horizontal orange bar. **Fig.3C:** Corresponding sliding-window analysis for human CTR9 (orange curve, same data as in Fig.3A), now showing also the sequence disorder profile (green curve). **Fig.3D:** PC profiles of the SPT6 IDR for different animal species are color-coded as indicated, wherein the PC profile for the human sequence essentially overlaps with those for the other sequences. The high-PC and high-disorder regions for the human IDR sequence are marked as in Fig.3B using orange and green bars and shaded areas; these colors do not share the meaning in the color code for the different sequences. **Fig.3E:** Left: bar chart of % identity from Clustal omega of SPT6 high-PC regions identified in different species relative to the human sequence. Right: % PC profile similarities (as defined in Materials and Methods) between the same high-PC regions in different species relative to the human sequence. **Fig.3F:** Same as Fig.3D but for CTR9. The high-PC and high-disorder regions for the human IDR sequence are marked as in Fig.3C. **Fig.3G:** Same as Fig. 3E but for CTR9. **Fig.3H:** Representation of the charge patterning and NCPR profile of high PC region from SPT6 from the indicated species. **Fig.3I:** Representative images of mCherry-tagged high PC regions of SPT6 from indicated species co-transfected with CFP-LacI-MED1^IDR^ (human) in the Lac array cell line. Top row shows mCherry fluorescence signal and bottom row shows CFP fluorescence signal. White box designates the Lac array focus using the CFP channel. Inset in top right corner of panels represents zoom in at CFP-LacI-MED1^IDR^ focus. Dashed white line designates the cell nucleus. Scale, 5μm. **Fig.3J:** Boxplot (min-max) of the partitioning as defined by the ratio of mCherry fluorescence signal inside/outside of CFP-LacI-MED1^IDR^ foci for SPT6 high PC regions from indicated species and an mCherry only control. Individual data points are plotted on top of the box plot. Dashed horizontal line indicates median of the control. *p*-values represent t-test for each high PC region relative to the control. Two-tailed unpaired t-tests relative to an mCherry control (N, 13) were performed for high PC regions of SPT6 from human (N, 12; p, <0.0001), mouse (N, 6; p, <0.0001), chicken (N, 8; p, <0.0001), frog (N, 8; p, 0.0003), or fish (N, 10; p, <0.0001). **Fig.3K:** Same as 3H but for CTR9 high PC regions from indicated species. **Fig.3L:** Same as 3I but for CTR9 high PC regions from indicated species. **Fig.3M:** Same as 3J but for CTR9 high PC regions from indicated species. Two-tailed unpaired t-tests relative to an mCherry control (N, 13) were performed for high PC regions of CTR9 from human (N, 9; p, <0.0001), mouse (N, 8; p, <0.0001), chicken (N, 11; p, <0.0001), frog (N, 8; p, <0.0001), or fish (N, 8; p, <0.0001).

We used this method to create PC profiles of CTR9 and SPT6, two proteins partitioned into MED1^IDR^ condensates and for which we have previously identified regions necessary and sufficient for partitioning^10^. Strikingly, the PC profile method identified regions within the N-terminal disordered region of SPT6 and the C-terminal disordered region of CTR9 (**Fig.3B and 3C**) which had been experimentally defined as necessary and sufficient for partitioning into CFP-LacI-MED1^IDR^ foci^10^. By overlaying disorder predictions from Metapredict^38,39^ (**Fig. 3B and 3C**, green line), plotted using the same average over sliding windows of 200 residues used for plotting the PC profile (orange line), we find that there is significant overlap between high PC and IDRs with high charge patterning (overlayed NCPR plot at the bottom). Interestingly, not all predicted disordered regions have high predicted PC (e.g. the C-terminal end of SPT6 in **Fig.3B**), as not all predicted disordered regions have the same charge or charge pattern characteristics. These results demonstrate the utility of the FH-RPA model in scanning full length proteins to identify regions with predicted high partitioning because those regions match experimentally validated regions responsible for partitioning of the protein.

### PC profiles are conserved despite a reduction in positional sequence conservation

To investigate whether PC profiles are an evolutionary conserved feature of proteins, we performed the PC profile analysis on SPT6 (**Fig.3D**) and CTR9 (**Fig.3F**) protein sequences from several animal species (human, mouse, chicken, frog and fish) (**Table S1**). The PC profiles of CTR9 or SPT6 from these different species were strikingly similar and all identified a high PC region in the same region of the protein (orange-shaded areas in **Fig.3D and 3F**). Sequence alignments of the high PC regions showed that % identity as defined by sequence alignment (Clustal omega) became lower relative to the human sequence with evolutionary distance for both SPT6 (**Fig.3E**, left) and CTR9 (**Fig.3G**, left). The poor conservation of positional sequence is expected for a region predicted to be disordered^40-43^. Nonetheless, when PC profiles of the high PC regions of SPT6 (**Fig. 3E**, right) or CTR9 (**Fig. 3G**, right) from different species were quantitatively compared to one another (Materials and Methods, Eq.10), they were nearly identical—as is also quite clear from inspection of **Fig.3D** and **3F**, suggesting that the high PC value is evolutionary conserved even though there is a reduction in positional sequence conservation. To test this prediction, we expressed the high PC regions of SPT6 (**Fig.3H-3J**) or CTR9 (**Fig.3K-3M**) from the indicated species and measured their partitioning into CFP-LacI-MED1^IDR^ foci in Lac array cells. While there was some targeting of these mCherry tagged proteins to other regions of the nucleus (**Fig. 3I and 3L**), high PC regions of SPT6 from different species (**Fig.3I**) and CTR9 from different species (**Fig.3L**) all partitioned into the CFP-LacI-MED1^IDR^ foci. Quantification focused on mCherry partitioning into CFP-LacI-MED1^IDR^ foci relative to an mCherry alone control showed that all predicted high PC regions from SPT6 (**Fig.3J**) and CTR9 (**Fig.3M**) have significant enrichment. These results show that the FH-RPA PC profile method can identify regions of full-length proteins that partition into CFP-LacI-MED1^IDR^, with PC profiles for SPT6 and CTR9 well conserved across species though the conservation of the amino acid residues along the sequence decreases significantly with evolutionary distance.

### FH-RPA model successfully predicts low-PC and high-PC IDR windows

Given the utility of FH-RPA to scan through arbitrary protein sequences and the striking agreement of predictions with experimental data, we next sought to scan through the proteome to identify novel sequences with high or low predicted PC values and test these predictions in the Lac array cell system. We decided to perform the sliding-window (tiling) analysis with 200 amino acid windows and focused this analysis on long (^3^ 200 amino acids) terminal disordered regions of nuclear proteins involved in transcription (**Fig.4A**).

**Fig. 4.**
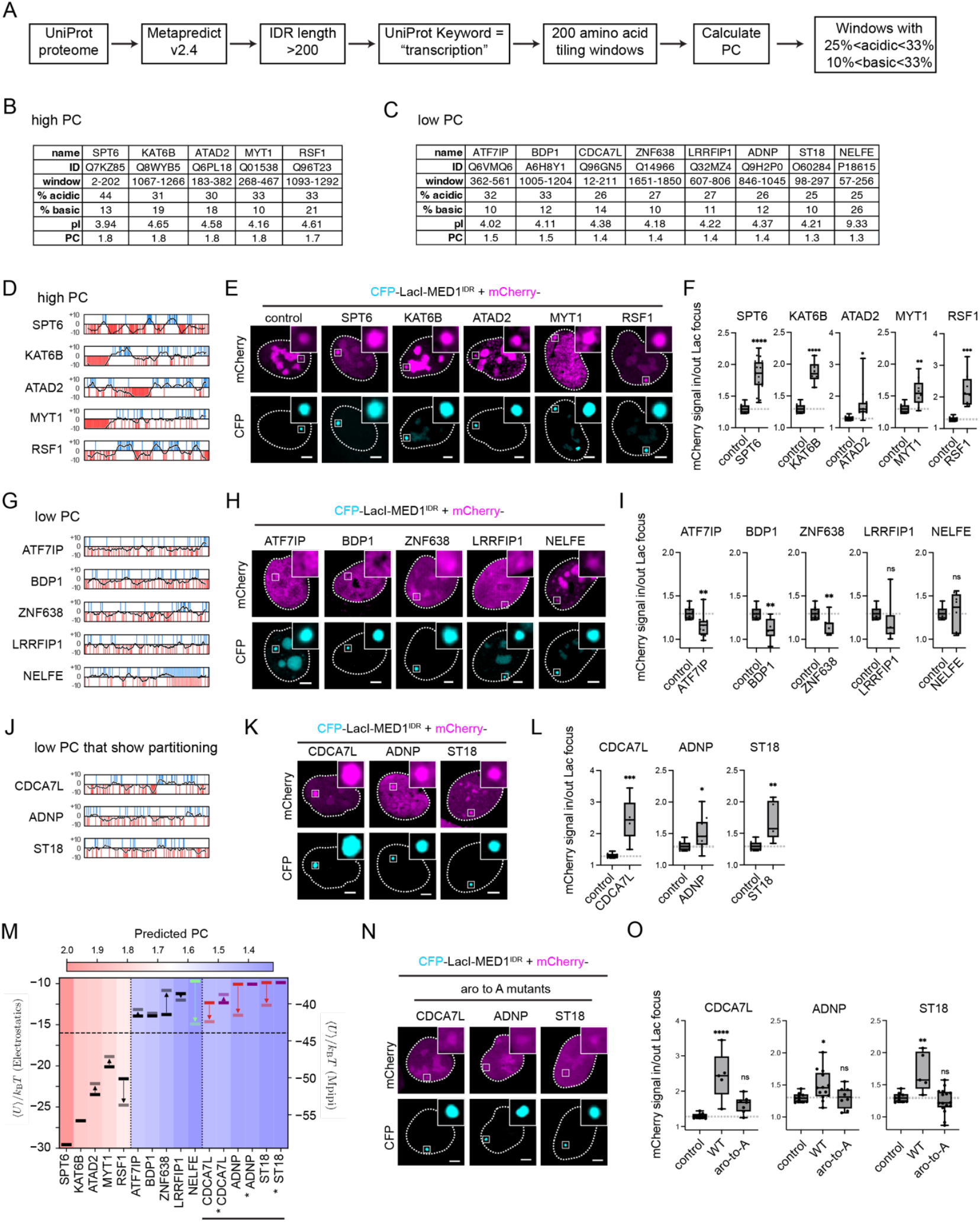
FH-RPA model successfully predicts low-PC and high-PC IDR windows from a large library of sequences. **Fig.4A:** Workflow to generate candidate set of 200 amino acid windows of IDRs from proteins involved in the same biological process (transcription) with comparable charge but either high or low predicted PC values. **Fig.4B:** Key parameters of the candidate set of 200 amino acid windows with predicted high PC values selected for subsequent experiments. Protein name and UniProt IDs and the amino acid boundaries (window) that define the region are provided. For a given region, % acidic and basic amino acids and isoelectric point (pI) as well as FH-RPA derived PC values (PC) are provided. **Fig.4C:** Same as Fig.4B but for the predicted low PC values. **Fig.4D:** Representation (as in Fig.2G) of the charge patterning and NCPR profile of 200 amino acid windows with predicted high PC. **Fig.4E:** Representative images of mCherry-tagged predicted high PC windows of indicated protein co-transfected with CFP-LacI-MED1^IDR^ in the Lac array cell line. Top row shows mCherry fluorescence signal and bottom row shows CFP fluorescence signal. White box designates the Lac array focus using the CFP channel. Inset in top right corner of panels represents zoom in at CFP-LacI-MED1^IDR^ focus. Dashed white line designates the cell nucleus. Scale, 5μm. **Fig.4F:** Boxplot (min-max) of the partitioning as defined by the ratio of mCherry fluorescence signal inside/outside of CFP-LacI-MED1^IDR^ foci for indicated 200 amino acid window and an mCherry only control. Individual data points are plotted on top of the box plot. Dashed horizontal line indicates median of the control. *p*-values represent t-test relative to the control. Two-tailed unpaired t-tests relative to an mCherry control (N, 9) were performed for 200 amino acid windows predicted to have high PC from SPT6 (N, 21; p, <0.0001), KAT6B (N, 6; p, <0.0001), ATAD2 (N, 13; p, 0.0181), MYT1 (N, 9; p, 0.0032), or RSF1 (N, 8; p, 0.0001). **Fig.4G:** Representation of the charge patterning and NCPR profile of 200 amino acid windows with predicted low PC. **Fig.4H:** Same as Fig.4E but for predicted low PC windows of indicated proteins. **Fig.4I:** Same as Fig.4F but for predicted low PC windows of indicated proteins. Two-tailed unpaired t-tests relative to an mCherry control (N, 9) were performed for 200 amino acid windows predicted to have low PC from ATF7IP (N, 11; p, 0.0083), BDP1 (N, 7; p, 0.0022), ZNF638 (N, 7; p, 0.0094), LRRFIP1 (N, 6; p, 0.2403), NEFLE (N, 8; p, 0.9475). **Fig.4J:** Representation of the charge patterning and NCPR profile of 200 amino acid windows with predicted low PC but which show anomalous partitioning into CFP-LacI-MED1^IDR^ foci. **Fig.4K:** Same as Fig.4E but for predicted low PC windows which show anomalous partitioning. **Fig.4L:** Same as Fig.4F but for predicted low PC windows which show anomalous partitioning. Two-tailed unpaired t-tests relative to an mCherry control (N, 9) were performed for 200 amino acid windows from CDCA7L (N, 5; p, 0.0003), ADNP (N, 13; p, 0.0345), or ST18 (N, 5; p, 0.0026). **Fig.4M:** Incorporation of non-electrostatic effects improve PC prediction. Sliding-sequence analysis is applied to IDRs with various RPA-predicted PCs (color coded). An intuitively chosen horizontal dashed line separates high (pink) and low (blue) predicted PCs. Boltzmann-weighted energies are averaged over both parallel and antiparallel alignments of sequence pairs (Fig.1B). Boltzmann-averaged energies computed using only screened Coulomb potentials as in Fig.1D (solid bars, left vertical scale) are compared to those computed using the Mpipi potentials^47^ (shaded bars, right vertical scale). Arrows indicate changes from screened Coulomb to Mpipi energies. Energies of IDRs with correctly predicted PC trends are shown in black, the 3 IDRs (in curly brackets) that were incorrectly predicted are in red, the one marginal case is in green, and the 3 aromatic to alanine mutants in Fig.4N,O are in magenta. **Fig.4N:** Representative images of mCherry-tagged predicted low PC windows with aromatic residues substituted for alanine (aro to A mutants) co-transfected with CFP-LacI-MED1^IDR^ in the Lac array cell line. Compare results to the data for the wildtype sequence presented in **Fig.4K**. Top row shows mCherry fluorescence signal and bottom row shows CFP fluorescence signal. White box designates the Lac array focus using the CFP channel. Inset in top right corner of panels represents zoom in at CFP-LacI-MED1^IDR^ focus. Dashed white line designates the cell nucleus. Scale, 5μm. **Fig.4O:** Boxplot (min-max) of the partitioning as defined by the ratio of mCherry fluorescence signal inside/outside of CFP-LacI-MED1^IDR^ foci for indicated 200 amino acid windows, the aromatic to alanine mutant (aro to A) and an mCherry only control. Individual data points are plotted on top of the box plot. Dashed horizontal line indicates median of the control. *p*-values represent one-way ANOVA with multiple comparisons tests vs. control. Ordinary one-way ANOVA with Dunnett’s multiple comparisons test to a “no IDR” mCherry control (N, 9) was performed for WT and aromatic to alanine substitution mutants (aro-to-A) for CDCA7L (WT: N, 5; p, <0.0001 and aro-to-A: N, 7; p, 0.1633), ADNP (WT: N, 13, p, 0.0499 and aro-to-A: N, 8; p, 0.9989), and ST18 (WT: N, 5; p, 0.0018 and aro-to-A: N, 16; p, 0.7019).

Starting with the UniProt reviewed human proteome, we first identified predicted disordered regions using Metapredict^38,39^, filtered for regions with ^3^ 200 amino acids, filtered for regions found within 10% of the N or C terminus, and finally filtered for disordered regions from proteins annotated with keyword “transcription” by UniProt. This yielded 865 IDRs with lengths ranging from 200 to 2869 amino acids. We then calculated FH-RPA derived PC values for 200 amino acid windows by scanning all sequences. This represented calculation of PC for 165,621 regions.

Given the contribution of electrostatics in the FH-RPA model calculation, we sought to compare 200 amino acid regions with comparable average charge but with different PC scores. We reasoned that comparing regions predicted to have different PC values but which differed dramatically in charge properties would not be a good test of the prediction. We therefore only focused on regions with relatively high acidic (25%<acidic<33%) and basic (10%<basic<33%) amino acid fraction and roughly similar isoelectric points (4.0 -4.65). This approach yielded a set of 4 regions predicted to have high PC and 7 predicted to have low PC. We included SPT6^IDR^ as our positive control and a 200 amino acid window of NELFE^IDR^ with the highest fraction of acidic amino acids as a negative control. Key parameters for SPT6^IDR^ and predicted high PC regions can be found in **Fig.4B** and those for NELFE and predicted low PC regions can be found in **Fig.4C**. All sequences can be found in **Table S1**. We expressed these sequences as mCherry fusions together with CFP-LacI-MED1^IDR^ in Lac array cells and measured their partitioning into CFP-LacI-MED1^IDR^ foci in cells. In this way, we were testing sequences of the same length, comparable average charge properties, but different predicted PCs.

NCPR profile analysis of the 4 new high PC regions showed that these regions had high local density of charge (**Fig.4D**). In Lac array cells, all four of these high PC regions partitioned into CFP-LacI-MED1^IDR^ foci (**Fig.4E**). While there was some variability of where these regions were targeted elsewhere in the nucleus (**Fig.4E**), quantification focused on mCherry enrichment at CFP-LacI-MED1^IDR^ foci shows significant enrichment for the four high PC regions relative to mCherry alone (**Fig.4F**). These results demonstrate that the FH-RPA can successfully identify well partitioned regions from an arbitrary list of sequences.

While four out of four (100%) of the high PC predictions were validated experimentally, only four out of seven (57%) of the low PC regions matched expectations and were not partitioned into CFP-LacI-MED1^IDR^ foci (**Fig.4G-4I**). Surprisingly, three out of seven were partitioned into CFP-LacI-MED1^IDR^ foci (**Fig.4J-4L**), suggesting that the FH-RPA model was missing some feature of these protein regions that were contributing to partitioning. There was nothing obvious about the difference in NCPR profiles (**Fig. 4G and 4J**) which prompted us to examine potential non-electrostatic interactions for these 3 outliers.

### FH-RPA model can be improved by including non-electrostatic effects

We examined possible reasons for why our currently implemented FH-RPA model predicted low PCs for regions of CDCA7L, ADNP, and ST18 that were shown experimentally to partition significantly into CFP-LacI-MED1^IDR^ foci (**Fig.4J-4L**). One limitation of our current FH-RPA model is that it does not account for non-electrostatic IDR-MED1^IDR^ interactions. We found one prominent difference in amino acid composition between the IDRs that are correctly predicted to have low PC (ATF7IP, BDP1, ZNF638, LRRHIP1) and the aforementioned three IDRs that are incorrectly predicted to have low PC: number of aromatic residues (F, Y, W). While the IDRs that are correctly predicted to have low PCs have at most 4 aromatic residues (ATF7IP: 4, BDP1: 1, ZNF638: 4, LRRHIP1: 2), the IDRs that are predicted to have low PCs but in fact have high or moderate experimental PCs have significantly more (≥10) aromatic residues (CDCA7L: 13, ADNP: 10, ST18: 10, NELFE: 10). Since MED1^IDR^ has a large number of cationic residues (75 K, 18 R), multivalent cation-p MED1^IDR^-IDR interactions could be playing a role^44^. It follows that a likely explanation for the inability of the current FH-RPA model to correctly predict the moderate to high experimental PCs of CDCA7L, ADNP, ST18 (and to a much lesser extent NELFE) is the model’s lack of consideration of non-electrostatic, and especially p-related interactions.

Aiming to explore whether incorporating p-related interactions can improve accuracy of PC prediction in general, we apply the sliding-sequence protocol in **Fig.1B,D** but now we compare the Boltzmann-averaged energies computed with only electrostatic (screened Coulomb) interactions against those computed with both electrostatic and non-electrostatic interactions. Several pairwise amino acid interaction schemes were developed recently to model IDR conformations and phase separations^45-48^. Here we adopt, as an example, the Mpipi potential^47^ because it has proven useful in accounting for p-related IDR interactions^24,47^. Because the non-electrostatic interactions are spatially short range, we use an inter-chain distance of 2*b* (which is smaller than those used in Fig.1C and Fig.1D) to ensure that these interactions are accounted for properly in the sliding-sequence analysis. The resulting changes in Boltzmann-averaged energies for the 16 IDR sequences studied in **Fig.4** are shown in **Fig.4M**. Assuringly, the IDRs predicted by FH-RPA to have high PCs (pink-shaded) have lower (more favorable) electrostatics-only Boltzmann-averaged energies (left vertical scale), as a group, than IDRs predicted by FH-RPA to have low PCs (blue-shaded), indicating that FH-RPA and the sliding-sequence protocol consistently capture the electrostatic effects on IDR partitioning into MED1^IDR^. When non-electrostatic effects are included, the Boltzmann-averaged energies are shifted (right vertical scale). The IDRs with high predicted PCs remain favorable (black and dark-gray bars in pink background), underscoring the robustness of the FH-RPA predictions for this group of sequences. The IDRs with correctly predicted low PCs remain similarly unfavorable or become even less favorable (black bars in blue background shifting to higher or only very slightly lower Boltzmann-averaged energies represented by the dark-gray bars), indicating that the FH-RPA predictions for this group of sequences are also robust. Interestingly, the group of sequences incorrectly predicted by FH-RPA to have low PCs (but in fact have moderate to high experimental PCs) all become more favorable (red and green bars shifting to lower Boltzmann-averaged energies), though not sufficiently favorable to match the favorability of those sequences correctly predicted by FH-RPA to have high PCs. Nonetheless, this exercise demonstrates that incorporation of non-electrostatic effects can improve accuracy of PC prediction. A possible first step in the future development of such an improved prediction scheme is to employ a mean-field approximation of non-electrostatic interactions^24^. Although such an approach does not account for the sequence pattern of uncharged residues, it may still offer advantages because of its account of hydrophobic and aromatic amino acid compositions (see SI for additional information).

To test this prediction from the FH-RPA + Mpipi correction, we mutated the aromatic residues in the experimental high-PC CDCA7L, ADNP, and ST18 IDR sequences to alanine. In **Fig.4M**, these aromatic to alanine mutant sequences are marked by an asterisk and their Boltzmann-averaged energies are represented by magenta bars, all of which remain high or become even slightly higher when non-electrostatic effects are included, indicating that partitioning is disfavored. Consistently, in Lac array cells, all three of the anomalous regions with aromatic to alanine substitutions had reduced partitioning into CFP-LacI-MED1^IDR^ foci compared to the wildtype sequence (**Fig.4N and 4O**). Quantification focused on mCherry enrichment at CFP-LacI-MED1^IDR^ foci showed that while the WT regions had significant enrichment relative to the mCherry-alone control, the aromatic to alanine substitutions reduced partitioning to levels comparable to or indistinguishable from the mCherry-alone control.

Taken together, these results demonstrate that multivalent contacts among disordered regions are a major contributor to the selective partitioning observed for condensates composed of the MED1^IDR^. The FH+RPA model allowed for calculating predicted PC values for large libraries of sequences. The predictions had striking qualitative agreement with previously published experimental results and predictions of the model were experimentally validated.

## Discussion

Partitioning of client molecules into biomolecular condensates is a general biophysical process for achieving microenvironments with regulated molecular compositions for biological functions^49^. For instance, earlier studies investigated the partitioning of small-molecule cancer therapeutics into nuclear condensates^50^ and highlighted partitioning of amphiphilic proteins as one of the means to assembling multilayered condensates^51^. Recent experimental^52^ as well as theoretical and computational^52-55^ investigations offered further physical insights into partitioning of small molecules into biomolecular condensates. These include the differential roles of hydrophobic and polar interactions^52,54^, utilization of simulated partitioning data to infer the general driving force for condensate assembly^55^, and the elucidation of a fundamental opposing effect against partitioning arising from the conformational entropy of the disordered protein scaffold^53^.

The present study focuses primarily on clients which are themselves disordered proteins. Disordered regions of proteins that serve as scaffolds of biomolecular condensates can promote the formation of biomolecular condensates that partition not only small molecules but also specific sets of proteins^10,12-14,31,32,56-59^, yet the mechanism underlying this specificity has been challenging to investigate. We recently demonstrated that the pattern of charged amino acids on IDRs was responsible for selective partitioning into condensates composed of MED1^IDR^, but we were unable to identify the mechanism of this specificity. Here we demonstrate that the selective partitioning of intrinsically disordered regions (IDRs) into MED1^IDR^ condensates can be largely explained by multivalent electrostatic interactions. Using a polymer physics-based approach, we developed the FH-RPA model, which integrates Flory-Huggins mean-field theory with a random phase approximation term to predict IDR partitioning based on sequence charge patterning. Our model successfully rationalized and predicted partitioning behaviors observed in previous experimental data and made predictions which were subsequently validated by experiments. By systematically scanning protein sequences, we identified novel high-partitioning IDRs and experimentally confirmed their behavior, highlighting the utility of this model in identifying regions responsible for partitioning. While our approach accurately predicted partitioning for most sequences, discrepancies for certain IDRs with significant aromatic content revealed the need to incorporate non-electrostatic interactions, such as cation-π interactions, to fully account for partitioning behavior. These results suggest that biomolecular condensate composition can, at least in part, be governed by statistical properties of disordered sequences.

Here we present theoretical effort focused largely on electrostatics, which plays important roles in the assembly^22,60^ and material properties^61^ of biomolecular condensates, and manifested also in salt-dependent phase separation of oligomeric peptides^62^ and biological IDRs^63-65^ as well as the significant impact of sequence charge pattern on the condensation of biological IDRs^60,66^ and the complex coacervation of multiple species of synthetic charged polymers^67,68^. Another exciting aspect of electrostatic contributions to condensate composition is the potential for regulation by post-translational modifications (PTMs). For example, multi-site phosphorylation has the potential to dramatically change the charge properties of disordered proteins. Indeed, the hyper-phosphorylation of the disordered C-terminal domain of RNA Polymerase II has been shown to change its partitioning behavior^56,57^. Future efforts to investigate whether PTMs regulate the partitioning of IDRs more generally will be an exciting future direction.

Taking a broader view, it should be noted that in the present implementation of our methodology, the empirical parameters of the model are determined by fitting only experimental data on (client) IDRs partitioning into MED1^IDR^ condensates. As such, the present implementation is not expected to address IDR partitioning into condensates scaffolded by other disordered proteins. Intuitively, how is the partitioning of a given client IDR into a MED1^IDR^ condensate related to the IDR’s partitioning into a condensate scaffolded by another disordered protein will depend on the nature of the other scaffold. For example, one may expect the degrees of client IDR partitioning will be similar if the sequence of the other scaffolding protein entails similar electrostatic interactions as those of MED1^IDR^, but the degrees of partitioning can be quite different if the interactions engaged by the other scaffolding protein are dominated by hydrophobic effects. The manner in which the present PC prediction method can be generalized to apply across condensates with different scaffolds remains to be investigated.

These limitations aside, our studies already highlight how the predicted PC values for MED1^IDR^-scaffolded condensates are conserved across different species even in the absence of positional sequence conservation. These observations add to the growing recognition of the evolutionary conservation of overall IDR properties such as charge, aromatic and hydrophobic contents, sequence motifs and repeats, and other molecular features that are sometimes not readily discernible by inspecting the primary amino acid sequences alone because sequence alignments often reveal little similarity; but these molecular features can be predictive of biological function^40,41,43^. For instance, conservation of function of sequentially highly diverged IDRs across species has been observed to be underpinned by conservation of simple electrostatic properties in the adaptor protein Ste50^69^ or binding affinities with a particular target in the case of p53TAD/MDM2^70^ without conservation of the primary amino acid sequence. The experimental observations in **Fig.3H-M** verify that the predicted high-PC windows for CTR9 and SPT6 indeed partition well into MED1 for IDRs from all five animal species we tested. Further efforts to investigate the co-evolution of IDR pairs may reveal additional features responsible for specificity.

The results described here demonstrate that multivalent contacts among disordered regions are capable of providing a physical account to explain the selective partitioning observed for condensates composed of the MED1^IDR^. Complementary to recent advances in applying forcefield parameters to predict IDR interaction properties^71,72^, including coarse-grained explicit-chain simulations of IDR partitioning into MED1-RNA condensates^73^, our high throughput analytical FH+RPA model allowed for efficient calculation of predicted PC values for large libraries of sequences. The predictions had striking qualitative agreement with previously published experimental results and predictions of the model were experimentally validated. In conclusion, our findings demonstrate, in the context of the experiments performed here, that selective partitioning can largely be explained by dynamic multivalent contacts among disordered regions without considering possible ordered-structure-mediated interactions.

## Acknowledgements

This work was supported by Canadian Institutes of Health Research (CIHR) grant NJT-155930, Natural Sciences and Engineering Research Council of Canada (NSERC) grants RGPIN-2018-04351 and RGPIN-2024-04167, and the computational resources provided by the Digital Research Alliance of Canada to H.S.C. and National Institutes of Health (NIH) grant GM147583 to BRS. This research was also supported in part by National Science Foundation grant NSF PHY-2309135 to the Kavli Institute for Theoretical Physics (KITP) and the Gordon and Betty Moore Foundation Grant No. 2919.02 through the KITP Program on “Physical Principles Shaping Biomolecular Condensates” in 2025.

## Author contributions

Conceptualization, JW, HSC, and BRS, data curation, JW, ND, and HL, formal analysis, JW and ND, funding acquisition, HSC and BRS, investigation, JW, ND, and HL, methodology, JW, ND, HL, and HSC, software, JW and ND, supervision, HSC and BRS, visualization, JW, ND, HSC, and BRS, writing – original draft, JW, HSC, and BRS, writing – review & editing, JW, ND, HL, HSC, and BRS

## MATERIALS AND METHODS

### A general screened electrostatic potential and the FH-RPA formulation

In general, the IDR-MED1 interaction energy *U*_*ij*_ (where the label *i* stands for the IDR in question and the label *j* stands for MED1) takes the usual screened-Coulomb form,

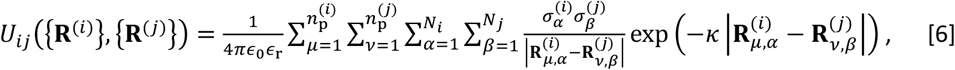

where *ϵ*_0_ and *ϵ*_r_ are, respectively, vacuum and relative permittivity,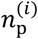 and 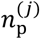 are, respectively, the numbers of chains of type *i* and type *j* in the system (chains labeled by *μ, v*), *N*_*i*_ and *N*_*j*_ are the numbers of residues in sequence of types *i* and *j* (residue positions labeled by *α*, β), 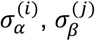 are, respectively, the charges at sequence positions *α*, β for sequence types *i, j*, 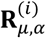 and 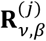 are the vectors for the spatial positions of (*i, μ, α*)- and (*j, v*, β)-labeled residues, and, as already defined, *κ* is the inverse Debye screening length.

Calculation of thermodynamic averages requires theoretically considering all possible chain configurations or conducting sufficient sampling thereof computationally. The intuitive “sliding-sequence’’ analysis (**Fig.1**) surmises that the limited set of configurations considered by sliding the IDR sequence over the MED1 sequence is representative of the full IDR-MED1 interaction such that it can provide semi-quantitative physical insights despite restricting to a particular {**R**^(*i*)^}, {**R**^(*j*)^} instead of covering all configurational possibilities rigorously. The results in **Fig.1C,D** were obtained using Bjerrum length *l*_*B*_ = *e*^2^/4*πϵ*_0_*ϵ*_r_*k*_B_*T* = 7 Å, where *e* is elementary (protonic) charge, *ϵ*_r_ ≈ 80 for bulk water (at *T* ≈ 300 K), and *κ*^-1^ = 10 Å corresponding to [NaCl] ≈ 100mM.

In the present FH-RPA formulation, we consider only electrostatic contributions to *χ*_*ij*_ (as indicated by the “e” subscript), viz.,

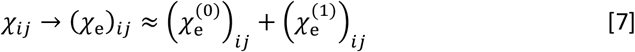

is composed of a zeroth order term

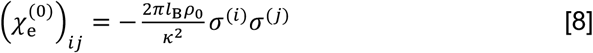

that depends only on the IDR and MED1 net charges *σ*^(*i*)^ and *σ*^(*j*)^, where 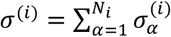 is the net charge of sequence type *i* (e.g., the net charges for MED1, NELFE, and SPT6 are +43, −2, and −62, and their chain lengths *N*_*i*_ or *N*_*j*_ are 626, 242, and 201, respectively). The first order term 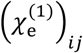 is given by Eq.2 in terms of

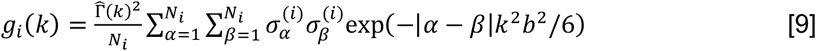

where *k* is reciprocal space (Fourier-transformed) coordinate variable, with small *k* associating with 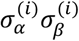 charge pairings with both large and small sequence separations (|*α* − *β*| s) and large *k* associating with charge pairings with only small sequence separations, and 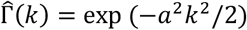 is a Gaussian smearing function (see SI for details). Software for computing *χ*_*ij*_ is publicly available from a Github webpage provided below.

For the NELFE and SPT6 examples in **Fig.2D** and **Fig.2E,F**, the theory-predicted PCs are overwhelmingly contributed by the first-order sequence charge pattern term, with 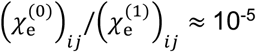 for NELFE and ≈ 10^−4^ for SPT6. An illustration of how the RPA term 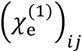 as defined above is affected by the IDR’s sequence charge pattern through the quantity *g*_*i*_(*k*) in our formulation is provided in **Fig.2D-F** for the NELFE and SPT6 examples.

### Calculating PC profiles

PC profiles are computed using 200-residue windows in the present study. For a given sequence, we determine 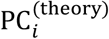 (Eq.1) for successive sequence windows comprising of 200 residues. A PC profile is a collection of such 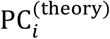 values, wherein the 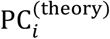 value of each 200-residue sequence window is plotted at the position of the 101th residue of the sequence windows (**Fig.3A**).

### Identifying high-PC regions

In the present study, we identify a high-PC region within a sequence by first locating the position of a 200-residue window with the maximum 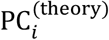 for the entire sequence. A high-PC region is a collection of consecutive 200-residue windows with 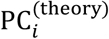 values of at least 90% of the maximum 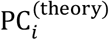 (**Fig.3A**).

### PC profile similarity score

Let 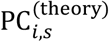 denote the theory-predicted partition coefficient (PC) of window *s* (Eq.1) of sequence *i* (consisting of segment of residues *s, s* + 1, …, *s* + *M* − 1 where *M* is the sequence window size). We then define 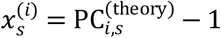 and 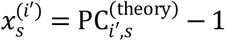 for two sequences of equal lengths *N*_*i*_ = *N*_*i*′_ ≡ *N*. A similarity score between their PC profiles may be constructed as

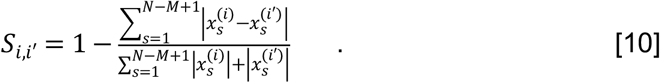

This similarity score is constructed such that 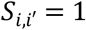 for two identical PC profiles whereas any sequence *i* has similarity 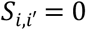 against the trivial profile 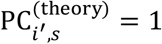 for all *s*. For sequences of different lengths, 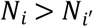, we define the similarity score 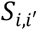 as the maximum 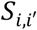 between *i*′ and all sub-sequences of *i* with length 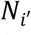, and vice versa if 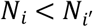. %PC similarity scores plotted in **Fig.3E** and **Fig.3G** is equal to 100 times 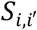.

### Cell Culture

All cells were grown, as previously described^12^, in full DMEM (Fisher Scientific 11995073) supplemented with 10% FBS, penicillin-streptomycin (Thermo Fisher 15140122) and GlutaMAX (Fisher Scientific 35050061) at 37 °C with 5% CO2 in a humidified sterile incubator.

### Processing coverslips of co-transfected cells

As previously described^12^, cells were seeded on coverslips (VWR 48366-067) in a 6-well plate and let adhere overnight. Transfections using Lipofectamine− 3000 Transfection Reagent with designated plasmids were incubated overnight. After cells were transfected and allowed to recover, cells were fixed in 4% paraformaldehyde (VWR BT140770) in PBS for 10 min at room temperature (RT) in the dark (plate wrapped with foil). After three washes in PBS for 10 min, samples were permeabilized with 0.5% triton X100 (Sigma T9284) in PBS for 10 min at RT. Then cells were incubated with 1:5000 Hoechst 33342 (Thermo Fisher, 62249) in Milli-Q water at RT in the dark (5 min). After washing once more with water, coverslips were mounted on slides (VWR 10144-820) with Vectashield (VWR 101098-042). Coverslips were sealed with nail polish (VWR 100491-940) and stored at 4°C.

### Microscopy

Images were acquired, as previously described^12^, using a Hamamatsu ORCA-Fusion C14440 digital camera and a CSU-W1 Yokogawa Spinning Disk Field Scanning Confocal System, and a 60x Plan Apo Lambda Oil Immersion objective (NA, 1.40). Thickness of Z-slices were 0.2 µm. Exposure time and laser intensity were the same for samples imaged in parallel. Laser lines used: 405nm, 488 nm, 561 nm, and 640 nm.

### Microscopy analysis

mCherry partitioning was calculated, as previously described^12^, by measuring the integrated fluorescence at the Lac array locus core as defined by fluorophore fused to LacI (highest intensity range on the Z-slice) and at a proximal spot in the nucleus (background) using FIJI 2D measurement tools and ROI tools. The same ROI area was used to measure the fluorescence at both background and focus. The quotient of these values (arbitrary units) was calculated as raw intensity at locus divided by raw intensity in the surrounding background.

### NCPR profile plots

Representation of the position of positive and negative charge along a protein sequence overlaid with net charge per residue (NCPR) profile provide a quick summary of the charge patterning found in a protein region. The position of acidic or basic amino acids are indicated as vertical red or blue lines, respectively. The horizontal black curve represents the NCPR profile using a 10 amino acid sliding window, wherein the net charge of successive 10-residue windows is plotted (10 times the net charge per residue). Consequently, each NCPR profile has a possible max of +10 and a possible min of -10 centered at 0 charge. X-axis is the length of the protein and y-axis is charge index.

### Statistics

GraphPad Software was used for statistical analysis. GraphPad notation for p values was used in figure panels: (ns p > 0.05, ^*^p ≤ 0.05, ^**^p ≤ 0.01, ^***^p ≤ 0.001, ^****^p ≤ 0.0001). For figure 2H and 2J, data were compiled from various experiments previously published^10^ to compare qualitative trends to the model’s predictions. For Fig.2H, ordinary one-way ANOVA with Dunnet’s multiple comparisons test to a “no IDR” mCherry control (N, 11) was performed for WT SPT6^IDR^ (N, 10; p, <0.0001), full scramble (N, 10; p, 0.6742), charge scramble (N, 10; p, >0.9999), and three independent non-charge scrambles (each with N, 10; p, <0.0001). For Fig.2J, ordinary one-way ANOVA with Dunnet’s multiple comparisons test to a “no IDR” mCherry control (N, 11) was performed for WT NELFE^IDR^ (N,10; p, 0.8805) and blocky (N, 10; p, <0.0001). For Fig.3J, two-tailed unpaired t-tests relative to an mCherry control (N, 13) were performed for high PC regions of SPT6 from human (N, 12; p, <0.0001), mouse (N, 6; p, <0.0001), chicken (N, 8; p, <0.0001), frog (N, 8; p, 0.0003), or fish (N, 10; p, <0.0001). For Fig.3M, two-tailed unpaired t-tests relative to an mCherry control (N, 13) were performed for high PC regions of CTR9 from human (N, 9; p, <0.0001), mouse (N, 8; p, <0.0001), chicken (N, 11; p, <0.0001), frog (N, 8; p, <0.0001), or fish (N, 8; p, <0.0001). For Fig.4F, two-tailed unpaired t-tests relative to an mCherry control (N, 9) were performed for 200 amino acid windows predicted to have high PC from SPT6 (N, 21; p, <0.0001), KAT6B (N, 6; p, <0.0001), ATAD2 (N, 13; p, 0.0181), MYT1 (N, 9; p, 0.0032), or RSF1 (N, 8; p, 0.0001). For Fig.4I and 4L, two-tailed unpaired t-tests relative to an mCherry control (N, 9) were performed for 200 amino acid windows predicted to have low PC from ATF7IP (N, 11; p, 0.0083), BDP1 (N, 7; p, 0.0022), ZNF638 (N, 7; p, 0.0094), LRRFIP1 (N, 6; p, 0.2403), NEFLE (N, 8; p, 0.9475), CDCA7L (N, 5; p, 0.0003), ADNP (N, 13; p, 0.0345), or ST18 (N, 5; p, 0.0026). For Fig.4O, ordinary one-way ANOVA with Dunnett’s multiple comparisons test to a “no IDR” mCherry control (N, 9) was performed for WT and aromatic to alanine substitution mutants (aro-to-A) for CDCA7L (WT: N, 5; p, <0.0001 and aro-to-A: N, 7; p, 0.1633), ADNP (WT: N, 13, p, 0.0499 and aro-to-A: N, 8; p, 0.9989), and ST18 (WT: N, 5; p, 0.0018 and aro-to-A: N, 16; p, 0.7019).

## Data Availability

The data supporting the findings of this study are available within the paper and its Supplementary Information. Should any raw data files be needed in another format they are available from the corresponding authors upon reasonable request.

## Software Availability

The effective Flory-Huggins parameter *χ*_*ij*_ in our FH-RPA formulation [Eq.1] were computed using our in-house, publicly available code EPIC-IDP (Effective Protein Interaction Calculator for Intrinsically Disordered Proteins) as a GitHub Python package at github.com/jwessen/Epic-IDP. A brief outline of the features in the package is also provided in SI.

