## Supplemental Information for "Sequence-based prediction of condensate composition reveals that specificity can emerge from multivalent interactions among disordered regions"

**July 31, 2025**

*for*

### Analytical Theory for Modeling Sequence-Dependent IDR Partitioning in MED1 Condensates

(*Reference numbers in square brackets refer to the list at the end of this SI document*)

**Random-Phase-Approximation (RPA) Theory for Binary Phase Separation of MED1 and a Charged IDR**

We first considered how sequence charge pattern affects IDR partitioning into MED1 condensate as a general question regarding how the sequences of a given IDR and that of MED1 (also an IDR) determine whether they phase separate together into a mixed condensed phase (i.e., a MED1 condensate is not presumed a priori); and if so, how the sequence of the IDR affects its concentrations in the condensed and dilute phases. With this aim, we applied the recently developed RPA theory for binary phase separation involving two solute components [1, 2] to systems comprising of MED1 and an IDR with a specific sequence as the two polymeric solutes. Here, as an exploratory calculation, only electrostatic interactions are taken into account aside from chain excluded volume. Accordingly, along the chain sequences of MED1 and the IDR, each arginine (R) or lysine (K) residue is assigned +1 electric charge (in units of protonic charge $e$), each aspartic (D) or glutamic (E) acid residue is assigned $-1$ charge, and all other residues are assigned zero charge. As detailed before [1, 2], the strength of electrostatic interactions in this class of model systems is dictated by the Bjerrum length $l_{\rm B} = e^2/4\pi\epsilon_0\epsilon_{\rm r}k_{\rm B}T$ and the Debye screening length $\kappa^{-1}$, where $\epsilon_0$ is vacuum permittivity, $\epsilon_{\rm r}$ is relative permittivity, $k_{\rm B}$ is Boltzmann's constant and $T$ is absolute temperature. The mathematical definition of this model, but generalized to multiple IDR species, is given by the formal development below [Eq. (S8)]. As an example, we applied this formulation to the natural (wildtype, WT) sequence of SPT6 and two of its variants, namely SPT6-IDR_charge_scramble (CS) and SPT6-IDR_scramble (S). With appropriate choices for $l_{\rm B}$ and $\kappa^{-1}$, RPA theory predicts binary phase separation for each of the three MED1-SPT6 systems (Fig. S1). Using a notation similar to that in previous studies [1, 2], the areas enclosed by the thick black curves in the phase diagrams in Fig. S1a–c are phase-separated regions. Dashed lines are tielines connecting co-existing phases with two different combinations of IDR and MED1 concentrations, from which a partition coefficient (PC) can be computed for the IDR, viz., PC = $[\text{IDR}]_{\text{condensed}}/[\text{IDR}]_{\text{dilute}}$, where $[\text{IDR}]_{\text{condensed}}$ and $[\text{IDR}]_{\text{dilute}}$ are, respectively, the IDR concentrations in the condensed (high-[IDR]) and dilute (low-[IDR]) phases at the two ends of a tieline for a given initial overall combination of [MED1] and [IDR].

Notably, Fig. S1d displays a good correlation between theoretically predicted and experimentally measured [3] PCs, supporting RPA binary phase separation theory as fundamentally capable of capturing the physical trend of charge-pattern-dependent partitioning of IDRs into MED1, thus arguing strongly for applying RPA generally to other IDR sequences. However, the PCs predicted in Fig. S1 from RPA binary phase separation theory are orders of magnitude too large in comparison with experimental values because of vanishing low IDR concentrations predicted for the dilute phase. A likely reason for this mismatch is the assumption that MED1/IDR phase separation is governed solely by the electrostatic interactions among MED1 and IDR chains (as an isolated system). But the physical reality is definitely much more complex, involving many yet unknown factors. Therefore, while recognizing the success of RPA in capturing how the behavioral trend of an IDR's partitioning depends on its sequence charge pattern, we construct below a more practical RPA formulation that offers predicted PC values that are comparable to those measured experimentally.

**A Simplified Flory-Huggins (FH) Approach to IDR Partitioning into a Preformed MED1 Condensate**

We now proceed to develop an approximate FH formalism for quantitative prediction of experimental PCs for IDRs into MED1 by presuming the pre-existence of a MED1 condensate while still leveraging RPA to account for the dependence on sequence charge patterns of IDR-MED1 interactions. Basically, this approach side-steps the difficulties in accurately understanding the driving forces for MED1 phase separation, especially in the complex intracellular environments in which some of the experimentally measured partitioning data were obtained. Moreover, by stipulating a preformed MED1 scaffold, the approach is computationally efficient because it does not entail constructing two-component phase diagrams to account for the phase separation of MED1, and thus allows for high-throughput scanning of large IDR sequence datasets to predict their PCs.

We begin the formalism development with the following consideration. In general, equilibrium partitioning of an IDR species $i$ into co-existing phases is governed by balancing its chemical potential $\mu_i^{(p)}$ in all phases labeled by $p$. To simplify the notation, all chemical potentials are given in units of thermal energy $k_{\rm B}T$ below. In general, one may split a given chemical potential into a translational entropy "ideal gas"-like contribution and an excess chemical potential $\mu_{{\rm ex.},i}^{(p)}$ originating from interactions, viz.,

$$\mu_i^{(p)} = \ln \rho_i^{(p)} + \mu_{{\rm ex.},i}^{(p)} \, , \tag{S1}$$

where $\rho_i^{(p)}$ is the number density (concentration) of $i$-type molecules in phase $p$. Population equilibrium between a dilute phase ($p$ = dil.) and a condensed phase ($p$ = cond.) implies that $\mu_i^{(\rm dil.)} = \mu_i^{(\rm cond.)}$. This equality can in turn be re-written as a relation between the partition coefficient ${\rm PC}_i \equiv \rho_i^{(\rm cond.)}/\rho_i^{(\rm dil.)}$ (defined as the ratio of concentrations in the two phases) with the difference in excess chemical potential between the two phases:

$${\rm PC}_i = \exp\left[\mu_{{\rm ex.},i}^{(\rm dil.)} - \mu_{{\rm ex.},i}^{(\rm cond.)}\right] \, . \tag{S2}$$

We now assume that all relevant interactions among the $M$ species ($i = 1, \ldots, M$) are well approximated by a Flory-Huggins (FH)-type quadratic free energy term based on Bragg-Williams random mixing,

$$f_{\rm int.} = -\frac{1}{\rho_0}\sum_{i=1}^{M}\sum_{j=1}^{M} \chi_{ij} N_i N_j \rho_i \rho_j \, , \tag{S3}$$

where $f_{\rm int.}$ is interaction free energy per unit volume (i.e., $f_{\rm int.}$ is free energy density) expressed in units of $k_{\rm B}T$. It follows that excess chemical potentials $\mu_{{\rm ex.},i} = \partial f_{\rm int.}/\partial \rho_i$ are linear in the densities of the molecular species,

$$\mu_{\text{ex.},i} = -\frac{2N_i}{\rho_0}\sum_{j=1}^{M}\chi_{ij}N_j\rho_j \ , \tag{S4}$$

where $N_i$ is the number of residues per $i$ chain, $\rho_0$ is a certain reference density (typically dominated by the solvent density) and $\chi_{ij}$ is an $M \times M$ symmetric matrix of dimensionless numbers that describe the pairwise interaction strengths between all molecular species in the system.

To make progress based on the above general consideration, we now make several simplifying assumptions regarding molecular densities in the respective phases. Suppose the condensed phase is dominated by one molecular species, e.g. $\rho_1^{(\text{cond.})} \gg \rho_{2,\ldots,M}^{(\text{cond.})}$ and the dilute phase is indeed dilute in all species, $\rho_i^{(\text{dil.})} \sim 0$. Under these conditions, we can approximate the excess chemical potentials as

$$\mu_{\text{ex.},i}^{(\text{dil.})} \sim 0 \quad , \quad \mu_{\text{ex.},i}^{(\text{cond.})} \approx -\frac{2N_1\rho_1^{(\text{cond.})}}{\rho_0}\chi_{i1}N_i \, , \tag{S5}$$

in the respective phases, where the first relation indicates that $\mu_{\text{ex.},i}^{(\text{dil.})}$ is small but nonzero. If we further assume that excess chemical potentials are small compared to the thermal energy $k_{\text{B}}T$, we can expect a linear relation between the partition coefficients $\text{PC}_i$ and the interaction parameters $\chi_{i1}$, viz.,

$$\text{PC}_i \approx c + \frac{2N_1\rho_1^{(\text{cond.})}}{\rho_0}\chi_{i1}N_i \, , \tag{S6}$$

where $c \approx 1$. In this approximation, the partitioning is solely determined by the interactions with the dominating species $i = 1$ of the condensate and the polymerization degrees $N_i$.

In what follows, we make the simplifying assumption that MED1 is the dominating IDR species ($i = 1$ in the above notation) acting as the scaffold of the condensate. We will use a polymer field theoretic framework to derive an approximate formula for $\chi_{i,\text{MED1}}$ determined by the amino acid sequences of MED1 and the partitioning IDR, with the latter acting as a client species (labeled by $i$). The numerical values of the model parameters will be determined by maximizing the Pearson correlation coefficient between a set of experimentally measured partition coefficients $\text{PC}_i^{(\text{exp.})}$ and the theoretical $\chi_{i,\text{MED1}}N_i$. Subsequently, we introduce two phenomenological parameters, $c_1$ and $c_2$, to arrive at a phenomenological formula for the IDR partition coefficients,

$$\text{PC}_i^{(\text{theory})} = c_1 + c_2\chi_{i,\text{MED1}}N_i \, , \tag{S7}$$

which is also given and discussed in the maintext. Determining the values of $c_1$ and $c_2$ by fitting $\text{PC}_i^{(\text{theory})}$ to $\text{PC}_i^{(\text{exp.})}$ then gives a predictive formula for partition coefficients entirely in terms of the IDR amino acid sequences. The coefficient $c_2$ encompasses unknown effects from solvent (including $\rho_0$) and condensed-phase MED1 concentration, while $c_1 \sim 1$ is expected from the above formal development for consistency and for a physically reasonable fit. To avoid introducing an additional fitting parameter, we do not consider the possibility of including another term linear in $N_i$ to account for a physically plausible change in client IDR configurational entropy upon partition into a condensed polymeric (MED1) environment [4].

**RPA-Derived FH Interaction Parameters for High-Throughput Prediction of Sequence-Dependent IDR Partitioning into Condensed MED1**

Proceeding to derive an effective FH $\chi_{ij}$ matrix using previous field-theoretic RPA modeling of IDR phase separation, our starting point is a general free energy per unit volume denoted by $f$ (in units of $k_{\rm B}T$) for $M$ IDR species with an RPA term accounting for electrostatic interactions through Gaussian fluctuations of the auxiliary field conjugate to charge density:

$$\begin{aligned}\frac{1}{\rho_0}f(\{\phi_i\}_{i=1}^{M}) =& \sum_{i=1}^{M}\frac{\phi_i}{N_i}\ln\phi_i + \phi_{\rm s}\ln\phi_{\rm s} - \sum_{i=1}^{M}\sum_{j=1}^{M}(\chi_{\rm h})_{ij}\,\phi_i\phi_j + \sum_{i=1}^{M}\sum_{j=1}^{M}\frac{2\pi l_{\rm B}\rho_0}{\kappa^2}\sigma_i\sigma_j\phi_i\phi_j \\ &+ \frac{1}{4\pi^2\rho_0}\int_0^\infty {\rm d}k\,k^2\ln\left[1+\frac{4\pi l_{\rm B}}{k^2+\kappa^2}\sum_{i=1}^{M}g_i(k)\rho_i\right].\end{aligned} \tag{S8}$$

This expression can be readily obtained by extending the framework developed in refs. [2, 5] to multiple IDR species with Debye screening. Here, the summation over $i, j = 1, \ldots, M$ run over all $M$ IDR species in the system. The other quantities in the above Eq. (S8) are:

- $N_i$: Number of residues (polymerization degree) of a species $i$ chain.
- $\rho_i$: Number density of species $i$, i.e. $\rho_i = n_i/V$ where $n_i$ is the number of $i$ chains and $V$ is the system volume.
- $\phi_i$: Volume fraction of species $i$, given by $\phi_i = N_i\rho_i/\rho_0$ where $\rho_0$ is the total bulk (overall) density.
- $\phi_{\rm s}$: Volume fraction occupied by solvent. This depends on the IDR densities through an imposed incompressibility condition $\phi_{\rm s} + \sum_{i=1}^{M}\phi_i = 1$.
- $l_{\rm B}$: Bjerrum length (controlling the strength of electrostatic interactions in a thermal environment).
- $\kappa$: Inverse Debye screening length. This is related to the ionic strength of the solution from implicitly treated ions. For a pure salt background, $\kappa$ is given by $\kappa = \sqrt{8\pi l_{\rm B}[{\rm NaCl}]}$ but other highly charged molecules not treated explicitly in this model (e.g. RNA) are expected to significanlty contribute to $\kappa$ in a biological setting.
- $\sigma_i$: Average residue charge (average net charge per residue) of an IDR of species $i$, i.e. $\sigma_i = \sum_{\alpha=1}^{N_i}\sigma_{i,\alpha}/N_i$ where $\sigma_{i,\alpha}$ is the electric charge of bead $\alpha$ on a species $i$ chain.

In the above Eq. (S8), the first two terms on the right hand side account for configurational entropies of the IDRs and solvent molecules. The third term represents a mean-field approximation of short-spatial-range interactions by using an effective FH $\chi_{ij}$ parameter wherein a subscript "h" alludes to hydrophobic-like effect though the formulation is generally applicable to any combination of different types of short-spatial-range effects including $\pi$-related interactions. This interaction term may be expressed as

$$(\chi_{\rm h})_{ij} = -\left(\int {\rm d}\mathbf{r}\, V_{\rm h}(|\mathbf{r}|)\right) \frac{\rho_0}{2N_iN_j} \sum_{\alpha=1}^{N_i}\sum_{\beta=1}^{N_j} \varepsilon_{r_\alpha^{(i)},r_\beta^{(j)}}\,, \tag{S9}$$

where $r_\alpha^{(i)}$ is the amino acid type of residue $\alpha$ on an IDR of species $i$. The matrix $\varepsilon_{r,r'}$ contains the pairwise-specific residue-residue interaction energies that multiplies an overall interaction potential $V_{\rm h}(|\mathbf{r}|)$, with $\varepsilon_{r,r'}V_{\rm h}(|\mathbf{r}|)$ in units of thermal energy $k_{\rm B}T$. The interaction parameters $\varepsilon_{r,r'}$ can be gleaned from existing parameter sets developed for coarse-grained molecular dynamic simulations of disordered proteins [6, 7, 8, 9, 10, 11]. In the present effort, however, we set $\chi_{\rm h} = 0$ to focus entirely on electrostatic interactions. Nonetheless, it is useful to note for future work that non-electrostatic contributions can be readily included through a $\chi_{\rm h}$ term.

The third term on the right hand side in Eq. (S8) corresponds to the mean-field theory (MFT) contribution from screened electrostatic interactions. This term is quadratic in the IDR densities and the coefficient in front of $\phi_i\phi_j$ defines a matrix of effective FH $\chi_{ij}$ parameters,

$$\left(\chi_{\rm e}^{(0)}\right)_{ij} = -\frac{2\pi l_{\rm B}\rho_0}{\kappa^2}\sigma_i\sigma_j\,. \tag{S10}$$

The final term in Eq. (S8) is the RPA contribution from electrostatic interactions. This term is an integral over fluctuation modes with wave numbers $k$. The quantity

$$g_i(k) = \hat{\Gamma}(k)^2 \sum_{\alpha=1}^{N_i}\sum_{\beta=1}^{N_i} \sigma_{i,\alpha}\sigma_{i,\beta}\,{\rm e}^{-|\alpha-\beta|k^2b^2/6} \tag{S11}$$

is a type of form-factor for the charge density of a single IDR species (labeled by $i$) modeled as a chain with Kuhn length $b$. $\hat{\Gamma}(k)$ describes the spatial distribution of charge for a single residue, which we take to be smeared by a Gaussian distribution, viz.,

$$\hat{\Gamma}(k) = {\rm e}^{-a^2k^2/2}\,. \tag{S12}$$

The smearing length $a$ thus serves as ultraviolet (UV) regulator since it eliminates the short wavelength modes ($k \lesssim a^{-1}$) in the RPA integral in Eq. (S8). Examples of $g_i(k)$ for several IDR-MED1 systems are provided in the maintext.

The RPA integral contains contributions to the free energy from all powers of $\rho_i$. For the purpose of this work, we are only interested in the quadratic term which can be viewed as a one-loop contribution to the effective $\chi_{ij}$ parameters quantifying the pairwise interaction strengths between the IDR species. Using the expansion $\ln(1+x) \approx x - x^2/2$ and dropping the physically inconsequential term linear in $\rho_i$ gives

$$\frac{1}{4\pi^2\rho_0}\int_0^\infty \mathrm{d}k\, k^2 \ln\left[1+\frac{4\pi l_\mathrm{B}}{k^2+\kappa^2}\sum_{i=1}^{M} g_i(k)\rho_i\right] \approx -\sum_{i,j}\left(\chi_\mathrm{e}^{(1)}\right)_{ij}\phi_i\phi_j\,, \tag{S13}$$

with

$$\left(\chi_\mathrm{e}^{(1)}\right)_{ij} = 2\pi l_\mathrm{B}^2\rho_0\int_0^\infty \mathrm{d}k\frac{k^2}{\left(k^2+\kappa^2\right)^2}\frac{g_i(k)}{N_i}\frac{g_j(k)}{N_j}\,. \tag{S14}$$

Note that Eqs. (S10), (S11), and (S14) are also provided and discussed in the maintext.

We use the effective $\chi_{ij}$-parameters to quantify the interaction strengths between IDRs in the present endeavor. In particular, we investigate to what extent the interaction strength between an IDR and MED1 controls its partitioning into MED1 condensates, as argued for in the previous section. Setting $\chi_\mathrm{h}=0$, we are left with a small number of model parameters $\{l_\mathrm{B},\kappa,\rho_0,a,b\}$ that enter into the calculation of the effective $\chi_{ij}$ parameter between two given sequences (i.e. given a pair of $\sigma_{\alpha,i}$). Without loss of generality, we can use the Kuhn length $b$ as our unit length such that it is no longer an independent tunable parameter. We quantify the correlation between measured partition coefficients and $\chi_{i,\mathrm{MED1}}N_i$ using Pearson's correlation coefficient $r$ which is invariant under shifts and rescaling. In other words, $\chi_{ij}N_j$ and $x+y\chi_{ij}Nj$ will give the same $r$ for any constants $x$ and $y$. This means that the total density, $\rho_0$, which only enters as an overall multiplicative factor to all contributions to $\chi_{ij}$ [Eqs. (S9), (S10) and (S14)] does not affect $r$ and can be omitted in the parameter fitting (because it can be absorbed into $y$). Similarly, any additive residue-independent contributions to $\chi_{ij}$ (modeling e.g., a common attraction between solvent and residues) will not affect $r$ since it can be absorbed into $x$.

We are thus left with three independent parameters $\{l_\mathrm{B},\kappa,a\}$ to be tuned to maximize the correlation with the experimental data to the following effective $\chi_{ij}$ parameter expression,

$$\chi_{ij} = -\frac{2\pi l_\mathrm{B}}{\kappa^2}\sigma_i\sigma_j + 2\pi l_\mathrm{B}^2\int_0^\infty \mathrm{d}k\frac{k^2}{\left(k^2+\kappa^2\right)^2}\frac{g_i(k)}{N_i}\frac{g_j(k)}{N_j}\,. \tag{S15}$$

As we will see in the following sections, the optimum is not a simple point in the $\{l_\mathrm{B},\kappa,a\}$ parameter space. Instead, there is an approximate degeneracy in the relevant region of parameter space where equivalent parameter points can be described as a trajectory in the $(l_\mathrm{B},\kappa)$-plane parametrized by the smearing length $a$, i.e., a curve $(l_\mathrm{B}(a),\kappa(a))$ in a range of $a$-values. This degeneracy implies that we effectively reduce the number of free parameters from three to two since the choice of $a$ is arbitrary within this $a$-range.

**Fitting Theory to Experimental IDR-MED1 Partitioning Data**

Among all the IDR sequences studied in the present investigation (Table S1), Table S2 provides the list of 30 IDRs (aside from MED1 itself) with experimentally determined partition coefficients (PCs) into MED1 (see maintext) that are used for the present parameter fitting. The residue contents of these IDR sequences are shown in Fig. S2. To assess the utility of RPA, it is instructive to first consider the non-RPA MFT contribution to the predicted PCs, namely $(\chi_\mathrm{e}^{(0)})_{ij}\propto-\sigma_i\sigma_j$ in Eq. (S10). At the MFT level of approximation, the correlation between experimental and MFT-predicted PCs is not influenced by any model parameters because it is solely governed by the net charges of the IDRs irrespective of their sequence charge patterns. The correlation turned out to be quite poor (Pearson correlation coefficient $r \sim 0.35$), indicating that net charge alone account for a little but not much of the physical chemistry of IDR partitioning into MED1 (see below).

Contrary to MFT, the RPA contribution given by Eq. (S14) is sensitive to the sequence charge pattern of the IDR and therefore is expected to better capture the sequence-dependent partitioning of IDRs into MED1. We now proceed to fit the experimental PC data to the expression in Eq. (S15) consisting of both the MFT and RPA contributions by optimizing a combination of $\{l_{\rm B}, \kappa, a\}$ values that maximizes the correlation between the PCs predicted by Eq. (S7) with $\chi_{i,{\rm MED1}}$ given by Eq. (S15). For terminological simplicity, quantities related to the $\chi_{ij}$ parameter in Eq. (S15) consisting of *both* the MFT and RPA contributions will be referred to as "RPA" quantities below unless specified otherwise.

An initial unconstrained numerical search in the $\{l_{\rm B}, \kappa, a\}$ parameter space did not converge to an optimal parameter point. Instead the optimization algorithm kept pursuing regions with increasing $l_{\rm B}$ until running into numerical problems related to the discretized integration of the RPA integral in Eq. (S15). Therefore, we ran a set of optimizations by imposing a maximal value of $l_{\rm B}$ evenly distributed on a logarithmic scale from $l_{\rm B}^{(\rm max)} = 0.1b$ to $l_{\rm B}^{(\rm max)} = 10^4 b$. The resulting optimal parameter point for each search are shown in Fig. S3. Interestingly, the correlation essentially saturates around $r \sim 0.67$ at $l_{\rm B}/b \gtrsim 10^1 - 10^2$, above which only a marginal increase in correlation happens even up to $l_{\rm B} \sim 10^4 b$. In this regime, where the parameters follow $\kappa^{-1} \lesssim a \ll l_{\rm B}$, there is an approximate degeneracy in the parameter space with respect to the correlation where the optimal points lie on a trajectory in the $(l_{\rm B}, \kappa)$-plane parametrized by the smearing length $a$.

**The Parameter Fitting Procedure**

To find parameter points that maximize the correlation between $\chi_{ij}$ and measured PC values, we performed several fits with varying upper cutoffs for $l_{\rm B}$. Each fit started out with 1,000 uniformly distributed points in the range $l_{\rm B} \in [0, l_{\rm B}^{(\rm max)}]$, $\kappa b \in [0, 10]$ and $a/b \in [0.01, 1]$. The algorithm then proceeds through 1,000 iterations where the best fit point was selected and 1,000 new points were generated in the vicinity of the best point. The points in Fig. S3 are the final points that yielded a Pearson correlation coefficient $r$ above 0.63 for the final point found in 200 independent runs with $l_{\rm B}^{(\rm max)}/b \in [0.1, 10^4]$ linearly spaced on a log-scale. Note that although the best-fit $l_{\rm B} \sim 10^4$ value (Fig. S4) is very high (corresponding to an extremely low temperature) for $r \sim 0.67$, the RPA-theory-experiment correlation achieved by using physically reasonable values of $l_{\rm B} = 2b$ and $\kappa = 0.1b^{-1}$ that match more closely with experimental conditions is only slightly weaker at $r = 0.647$ (with $a = 0.15b$ and $b = 3.8$ Å is the virtual $C_\alpha$–$C_\alpha$ bond length). This result suggests convincingly that our RPA formulation is capable of capturing the essential physics of experimental IDR-MED1 partitioning. With this recognition in mind, we will use the optimized parameters for $r \sim 0.67$ for moderately enhanced numerical accuracy.

The MFT- and RPA-predicted PCs are compared in Fig. S4. The upper panel (Fig. S4a) depicts the aforementioned weak correlation between experimental and MFT-predicted PCs with $r = 0.346$. Here, among the strongest outliers in the scatter plot are CTR9 and

NELFE-tentacle_blocky which are blocky polyampholytes with a high fraction of charged residues but a small net charge. The lower panel (Fig. S4b) showcases the correlation between experimental PCs and RPA-predicted PCs using the best-fit parameter points in Fig. S3, exhibiting a significantly improved correlation of $r = 0.671$ relative to the MFT only case. Of particular interest is the fact that IDRs with few charged residues (FUS and EWSR1) are now correctly predicted to have small partitioning into MED1. Nonetheless, the correlation is far from perfect: the predicted PCs for the blocky polyampholytes CTR9 and NELFE-tentacle_blocky still significantly underestimate the experimental values, though there are small yet discernible improvements vis-à-vis MFT even for these two outliers. It is also noteworthy that the best-fit values for $c_1$ and $c_2$ in Eq. (S7) for MFT are $c_1 = 1.45$, $c_2 = 1.72 \times 10^{-4}$ and for RPA are $c_1 = 0.943$, $c_2 = 7.73 \times 10^{-8}$, consistent with the $c_1 \approx 1$ assumption used in our formal derivation.

To assess the robustness of these fits, we performed bootstrap analyses by repeating the calculation of the Pearson correlation coefficient 500,000 times, each time using 19 randomly selected points (with replacements) among the 19 wildtype client IDR sequences in Table S2 (not including MED1 itself), yielding a bootstrap distribution $P(r)$ of correlation coefficients $r$ (Fig. S5). The $Z$-scores, defined as $Z = (r - \langle r \rangle)/\sigma_r$, where $\langle r \rangle$ and $\sigma_r$ are the mean and standard deviations of the bootstrap distributions, are $Z \approx -0.25$ for RPA and $Z \approx -0.28$ for MFT. The similarity in value between $r$ and $\langle r \rangle$ and the relatively small $\sigma_r$ for RPA shown in Fig. S5 indicate that the $r \approx 0.67$–$0.70$ correlation from our RPA fit (Fig. 2C in the maintext and Fig. S4) is typical (not an outlier) of the experimental-theoretical PC sample in Table S2.

**Predicting MED1 Partition Coefficients for Extensive IDR Sequence Databases**

We used the best-fit set of parameters discussed above and stated in the caption for Fig. S4 to calculate the RPA-predicted PCs for all sequences in the Metapredict v2.4 IDRome [12, 13] with sequence lengths shorter than $N = 4{,}100$ amino acid residues. The distribution of predicted PC values is provided in Fig. S6.

To gain further insight, we explored how the RPA-predicted PC values are affected by the composition (content) of charged residues in the IDR sequences, as follows. For a sequence with $N$ residues, let $N_+$ and $N_-$ denote, respectively, the number of positively and negatively charged residues, then $\text{FCR} \equiv (N_+ + N_-)/N$ is the fraction of charged residues, and $\text{NCCR} \equiv (N_+ - N_-)/(N_+ + N_-)$ is the net charge per charged residue. Using these characterizations as discriminants, we classify IDR sequences into four categories:

- $< 5\%$ *charged residues:* Sequences with FCR $< 0.05$.
- *Polyampholytes:* Sequences with FCR $> 0.2$ and NCCR $< 0.2$.
- *Poly-cations:* Sequences with FCR $> 0.2$ and NCCR $> 0.8$.
- *Poly-anions:* Sequences with FCR $> 0.2$ and NCCR $< -0.8$.

Fig. S7 shows the distribution of RPA-predicted PC values among the categorized sequences. The scatter plot in Fig. S8 illustrates further the relationship between the predicted PC values and the average charge per residue $\bar{\sigma} = (N_+ - N_-)/N$ of the IDR sequences. Both

Fig. S7 and Fig. S8 indicate that the largest variation in RPA-predicted PCs is observed among polyampholytic IDRs.

Notably, the RPA-predicted PC values for the databases reveal a significant positive correlation between sequence length $N$ and predicted PC (Fig. S9). This trend is not unexpected because longer IDRs tend to entail stronger interactions per IDR molecule with any scaffold molecule such as MED1 simply because there are more amino acid residues along the IDR chain to engage in possibly favorable interactions. Aiming to decipher the basic relation between PC and sequence charge pattern here, we factor out $N$-dependence by further focusing on the predicted PCs of all 200-residue windows (i.e., with the same chain length) in the Metapredict IDRome sequences. As described in the maintext, 200-residue windows with similar net charges from 11 different proteins are selected for experimental testing together with SPT6$^{\rm IDR}$ and a 200-residue window of NELFE$^{\rm IDR}$ as experimental controls. The comparison between experimental data and theoretical predictions is reported in the maintext.

Though not pursued in the present study, we note that the theoretical framework developed here may also be harnessed to discover novel charge-scrambled IDR sequences that have elevated or attenuated partitioning compared to that of the wildtype sequence. For example, one may proceed by using a genetic algorithm to search the sequence space defined by possible re-arrangements of the order of amino acid residues along the chain sequence, i.e., by residue position swapping/scrambling that maintain the same overall amino acid residue composition [14].

#### Available Software

The effective $\chi_{ij}$ in Eq. (S15) were computed using our in-house code EPIC-IDP (Effective Protein Interaction Calculator for Intrinsically Disordered Proteins). This code is publicly available as a Python package on GitHub at `github.com/jwessen/Epic-IDP`.

To use the package, one first defines a `chi_effective_calculator` instance which takes interaction parameter values as input, for example:

```
from epic_idp import chi_effective_calculator

cec = chi_effective_calculator( rho0  = 1.                ,\
                                lB    = 2.                ,\
                                kappa = 0.1               ,\
                                a     = 0.15              ,\
                                Vh0   = 0                 ,\
                                interaction_matrix = 'KH-D' )
```

The arguments correspond to

- `rho0` $\leftrightarrow \rho_0 b^3$
- `lB` $\leftrightarrow l_{\rm B}/b$
- `kappa` $\leftrightarrow \kappa b$
- `a` $\leftrightarrow a$
- `Vh0` $\leftrightarrow \int d\mathbf{r} V_{\rm h}(|\mathbf{r}|)$

The `interaction_matrix` argument refers to the data set (hydrophobicity scales or interaction matrices) used for contact energies $\varepsilon_{r,r'}$. It has to be one of the following:

- `'KH-D'` (S3 Data in [6])
- `'Mpipi'` (using the 20-by-20 matrix for amino-acid pairs [7])
- `'Mpipi_RNA'` (Using the 24-by-24 matrix including RNA bases in [7])
- `'CALVADOS1'` (Using the original CALVADOS dataset [8])
- `'CALVADOS2'` (Using the updated CALVADOS dataset [9])
- `'HPS'` (Table S1 in [6])
- `'URRY'` (Table S2 in [10])
- `'FB'` (Table S7 in [11])

After creating the `chi_effective_calculator` instance, we define which IDR sequences we are interested in. The IDR sequences are added sequentially with no upper limit on how many sequences can be added. For example:

```
# Sequences in FASTA format
sequences = [ 'EHHSGSQGPLLTTGDLGKEKTQ' ,
    'RKQDEEERELRAKQEQEKELLRQKLLKEQEEK' , 'AGREAKRR' ]

# Names of the IDPs
names = [ 'seq1' , 'seq2' , 'seq3' ]

for seq, name in zip(sequences,names):
    cec.add_IDP(name, seq)
```

When adding an IDR sequence, the program directly computes the double sum over residues in $g_i(k)$ (as defined in Eq. (S11)) which constitutes the most computationally heavy step of the effective $\chi_{ij}$ parameter calculations. After all IDRs are added, the $\chi_{ij}$ elements (with an overall factor $\rho_0$) are computed using the functions of the `chi_effective_calculator` instance, for example:

```
# Returns the effective chi-parameter for the seq1-seq2 pair
cec.calc_chi_eff('seq1', 'seq2')

# Returns the full M-by-M matrix of chi parameter (M is the
    number of added IDPs)
cec.calc_all_chi_eff()
```

Full documentation of the EPIC-IDP module can be found in the `readme.md` file at `https://github.com/jwessen/Epic-IDP`.

**Table S1**: This table is provided separately as an Excel (`xlsx`) file.

**Table S2**: All IDR sequences used for fitting the PC-predicting parameters. Listed under PC (experiment), PC (MFT), and PC (RPA) are, respectively, the PC values measured experimentally, predicted by mean field theory (MFT) by considering only the net charges of the IDRs, and predicted by the RPA formulation that takes into account the sequence charge patterns of the IDRs. Experimental PCs are averaged over 10 measurements, corresponding standard deviations are provided after the ± signs. Listed IDR sequences from MED1 to RCOR3 are natural (wildtype) sequences. Sequences starting with Synthetic_IDR_1 and below are synthetic variants; their abbreviated names used for the labels in Fig. S4 are given in parentheses. An `xlsx` version of this table (`Table_S2.xlsx`) is also provided separately.

| IDR Name (abbrev. in Fig. S4) | PC (experiment) | PC (MFT) | PC (RPA) |
|---|---|---|---|
| MED1 | 1.73±0.49 | 1.23 | 1.65 |
| CTR9 | 2.74±0.84 | 1.48 | 1.87 |
| SPT6 | 2.10±0.42 | 1.75 | 2.00 |
| IWS1 | 1.76±0.35 | 1.91 | 2.21 |
| NELFA | 1.39±0.16 | 1.39 | 1.30 |
| NELFE | 1.15±0.11 | 1.46 | 1.33 |
| HP1a | 1.10±0.22 | 1.41 | 1.07 |
| LEO1 | 1.53±0.28 | 1.91 | 2.08 |
| PAF1 | 1.52±0.21 | 1.67 | 1.56 |
| DDX4 | 1.42±0.19 | 1.47 | 1.28 |
| EWSR1 | 0.92±0.11 | 1.47 | 0.98 |
| EP300 | 0.94±0.20 | 1.37 | 1.14 |
| FUS | 1.10±0.13 | 1.47 | 0.97 |
| Ki67_R12_pm9x2 (Ki67) | 1.14±0.12 | 1.49 | 1.39 |
| NICD | 1.20±0.21 | 1.55 | 1.22 |
| NPM1 | 1.37±0.20 | 1.54 | 1.50 |
| MeCP2 | 0.88±0.17 | 1.28 | 1.51 |
| CBX2 | 1.27±0.18 | 1.29 | 1.55 |
| KDM1A | 1.10±0.10 | 1.49 | 1.22 |
| RCOR3 | 1.29±0.18 | 1.45 | 1.06 |
| Synthetic_IDR_1 (Synthetic_1) | 1.76±0.13 | 1.69 | 1.80 |
| Synthetic_IDR_2 (Synthetic_2) | 1.77±0.22 | 1.66 | 1.75 |
| Synthetic_IDR_3 (Synthetic_3) | 2.29±0.24 | 1.67 | 1.76 |
| Synthetic_IDR_4 (Synthetic_4) | 1.78±0.33 | 1.66 | 1.74 |
| Synthetic_IDR_5 (Synthetic_5) | 1.80±0.26 | 1.66 | 1.71 |
| SPT6-IDR_scramble (SPT6_S) | 1.08±0.08 | 1.75 | 1.67 |
| SPT6-IDR_charge_scramble (SPT6_CS) | 1.24±0.16 | 1.75 | 1.67 |
| SPT6-IDR_noncharge_scramble_1 (SPT6_NCS_1) | 2.19±0.42 | 1.75 | 2.00 |
| SPT6-IDR_noncharge_scramble_2 (SPT6_NCS_2) | 2.01±0.49 | 1.75 | 2.00 |
| SPT6-IDR_noncharge_scramble_3 (SPT6_NCS_3) | 2.20±0.42 | 1.75 | 2.00 |
| NELFE-tentacle_blocky (NELFE_blocky) | 2.85±1.16 | 1.46 | 1.62 |

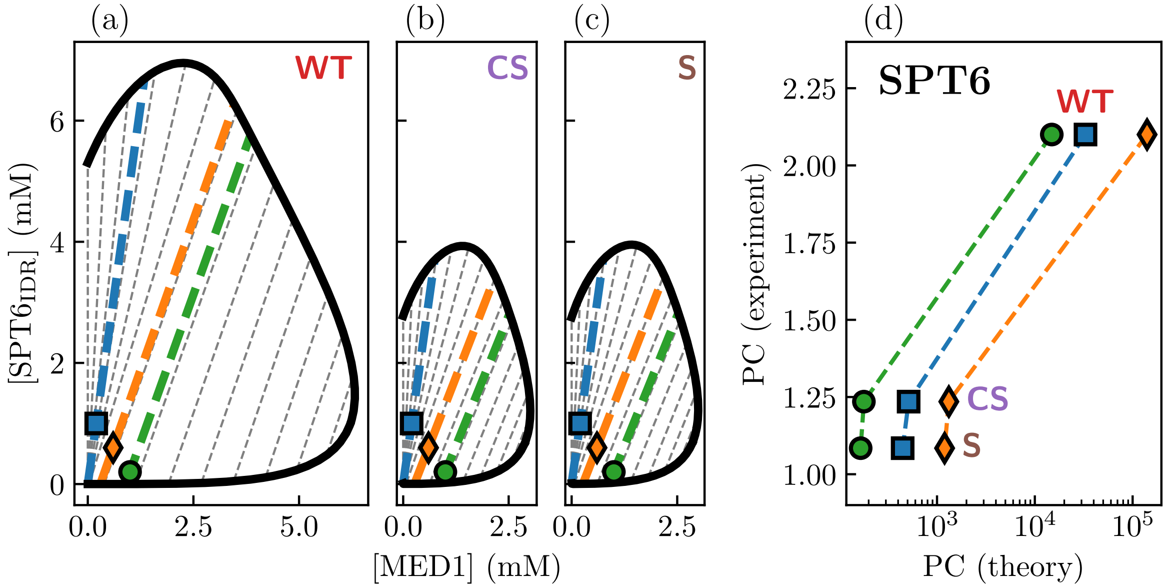


**Figure S1**: Binary phase separation predicted by random phase approximation (RPA) for MED1 and SPT6 variants. Each SPT6 IDR sequence [(a) WT, (b) CS, and (c) S, see Tables S1 and S2] is paired with MED1, for which a two-component binodal (co-existence curve, solid black) is obtained from RPA phase separation theory [1, 2]. To avoid dealing with potentially more complicated situations involving ternary phase separation in this exploratory calculation, a Bjerrum length $l_{\mathrm{B}} = 0.5b$ where $b$ is the Kuhn length in the RPA theory, a total solvent + MED1 + IDR bead density $= 5b^{-3}$, and a Debye screening length $\kappa^{-1}$ equivalent to [NaCl] = 500 mM are used to obtain the phase diagrams in (a–c). Black and color dashed lines in (a–c) are tielines. Three different combinations of initial (overall) concentrations that are inside all sequence pairs' co-existing regions are compared: ([SPT6$_{\mathrm{IDR}}$],[MED1])/mM = (0.2,1.0), (0.6,0.6), and (1.0,0.2). (d) Correlation between experimentally measure partition coefficients (PCs) [3] and theoretical PCs computed from the [SPT6$_{\mathrm{IDR}}$] values at the two ends of the color tielines in (a–c) as described in the text. Dashed color lines highlighting the trends in (d) are merely a guide for the eye.

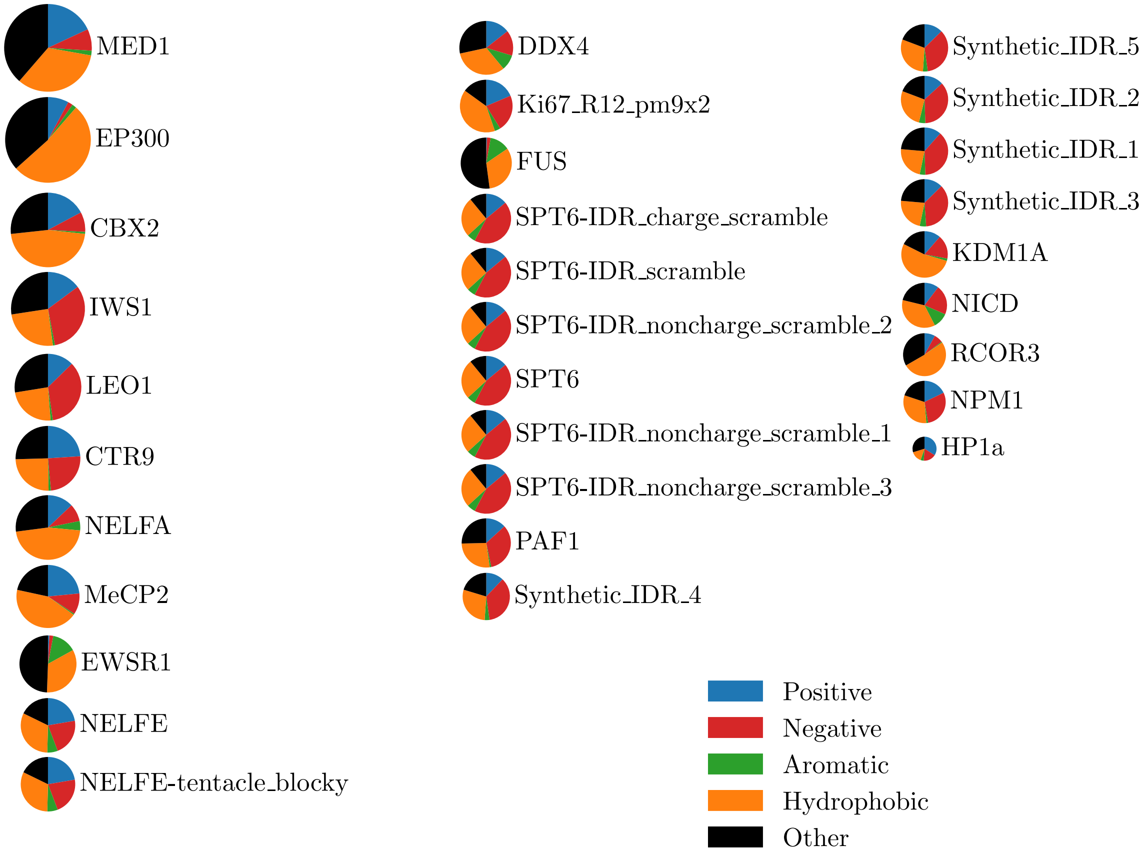


**Figure S2**: Amino acid contents (compositions) of every IDR in Table S2 considered for parameter fitting. The area of the circles is proportional to the number of amino acid residues per chain, ranging from $N = 626$ for MED1 to $N = 47$ for HP1a. Notably, FUS and EWSR1 have large content of aromatic residues but few charged residues. Here, in terms of the one-letter code for amino acids, positive residues are R, K; negative residues are D, E; aromatic residues are F, W, Y; hydrophobic residues are A, I, L, M, V, G, P; and the remaining residue types are classified as “other”.

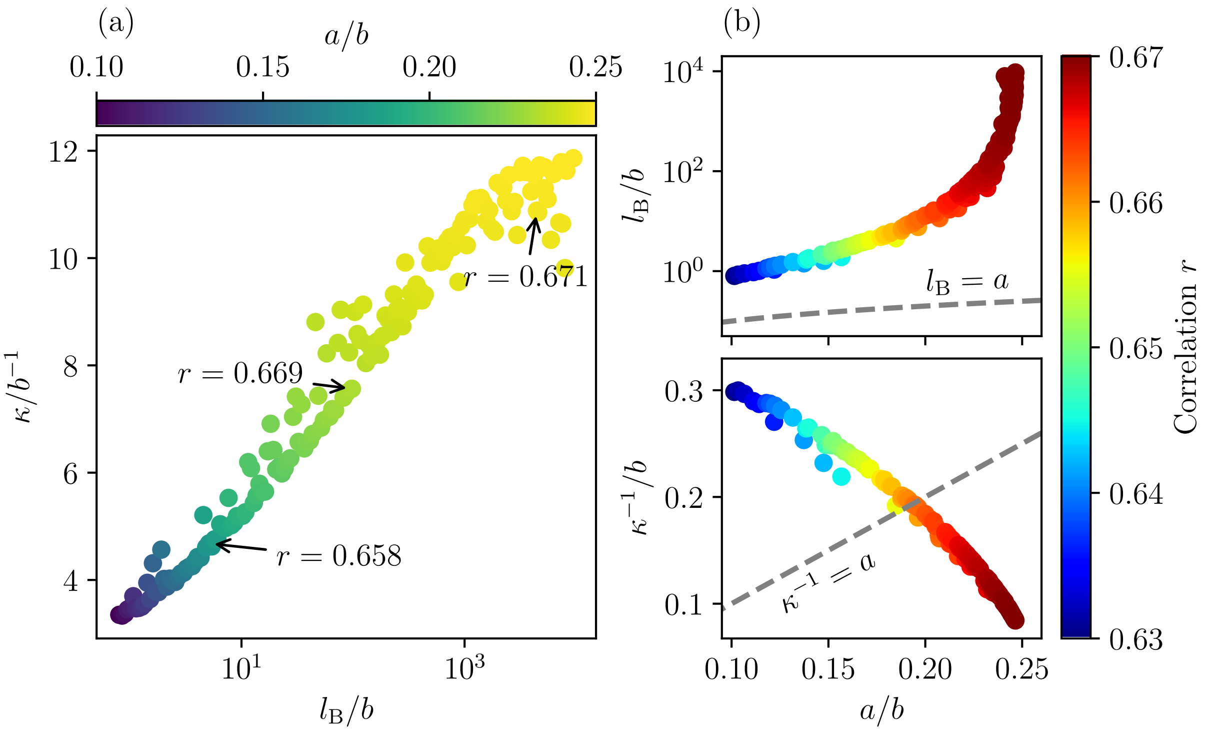


**Figure S3**: Results of the parameter space scans for optimizing Pearson's correlation coefficient $r$ against experimentally measured partition coefficients. Each data point is the result of a separate optimization run with a different imposed maximal value of $l_{\mathrm{B}}$. (a) Parameters optimizing the correlation at different $l_{\mathrm{B}}^{(\mathrm{max})}$. (b) "Flow" of optimized $l_{\mathrm{B}}$ and $\kappa^{-1}$ parameters as smearing length $a$ is varied as a fitting parameter.

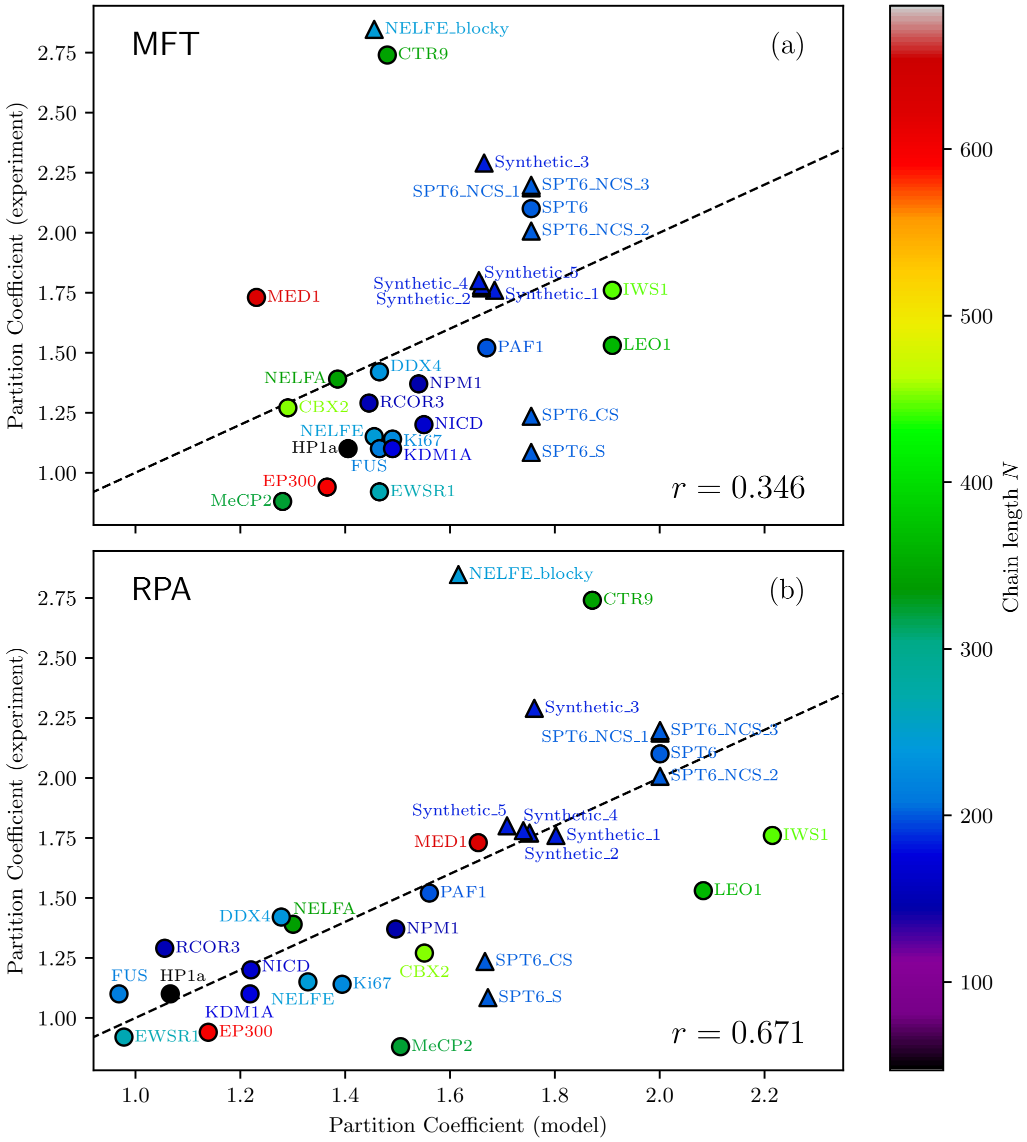


**Figure S4**: Optimized predictions of experimental partition coefficients (PCs) by theory. Experimental data on PCs for all IDR sequences into MED1 as listed in Table S2 are used in the fits. Natural (wildtype, WT) and synthetic IDRs are represented, respectively, by circles and triangles in the scatter plots. The chain lengths of the IDRs are colored coded by the scale on the right. The dashed ($x = y$) lines mark the hypothetical situation when model/theoretical PCs match perfectly with experimental PCs. (a) MFT prediction of MED1-IDR partitioning takes into account only the composition of amino acid residue contents of IDRs but not their sequence pattern. Here, strongest outliers are the blockly polyampholytes CTR9 and NELFE_blocky. The correlation is quite weak with Pearson correlation coefficient $r \approx 0.35$. (b) RPA prediction takes into account sequence charge patterns by incorporating the RPA-derived effective $\chi_{ij}$ parameter in Eq. (S15) using the optimized parameter set $\{l_B, \kappa, a\} = \{9437.9b, 11.861b^{-1}, 0.24651b\}$. This leads to a significant improvement in correlation between experimental and model PCs with $r \approx 0.67$, though a few outliers persist.

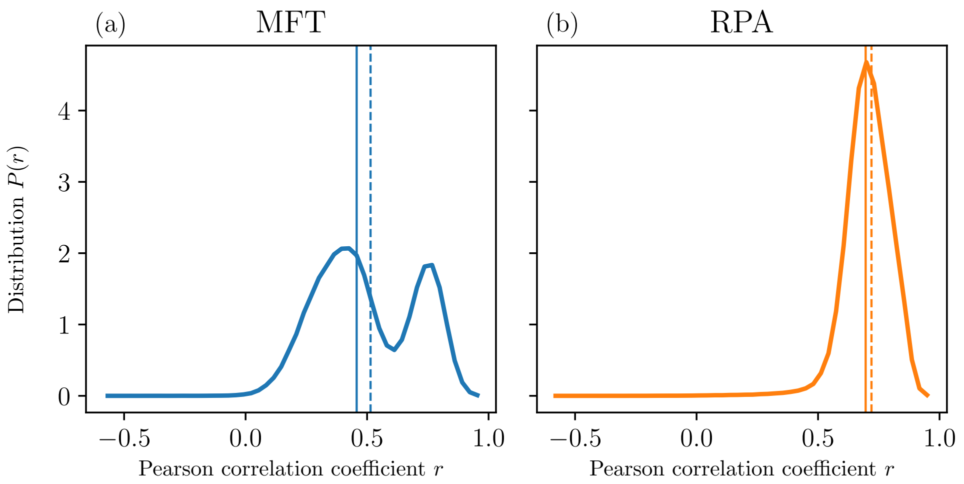


**Figure S5**: Bootstrap analysis of experimental-theoretical PC fits. The distribution $P(r)$ of Pearson correlation coefficient $r$ (continuous curve) over 500,000 samples each with 19 randomly chosen data points (with replacements) for the 19 wildtype sequences from Table S2 (same sequences considered in Fig. 2B,C in the main text, MED1 itself is not included) is shown separately for mean-field theory (MFT, left panel) and random phase approximation (RPA, right panel). The correlation coefficient $r$ for the overall fit (using each data point only once without replacement) and the average correlation coefficient $\langle r \rangle$ are marked, respectively, by the vertical solid and dashed lines.

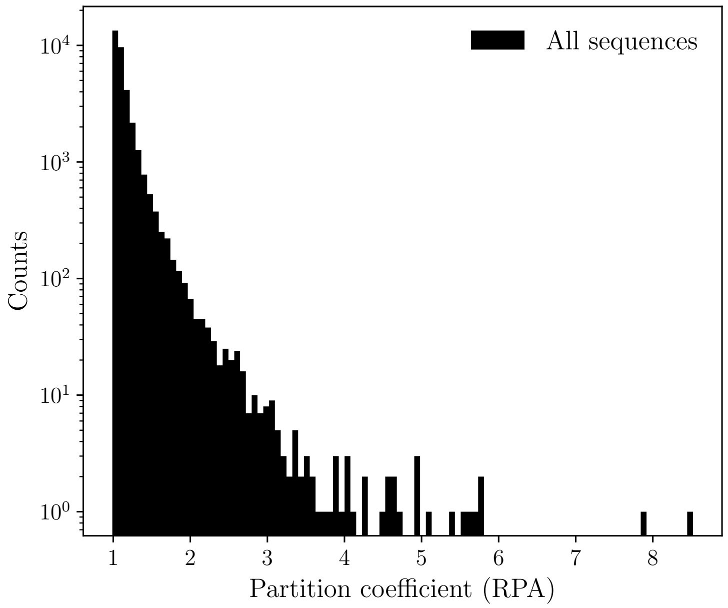


**Figure S6**: Distribution of RPA-predicted partition coefficients for all sequences in the Metapredict v2.4 IDRome with chain lengths (number of amino acid residues) $N < 4{,}100$.

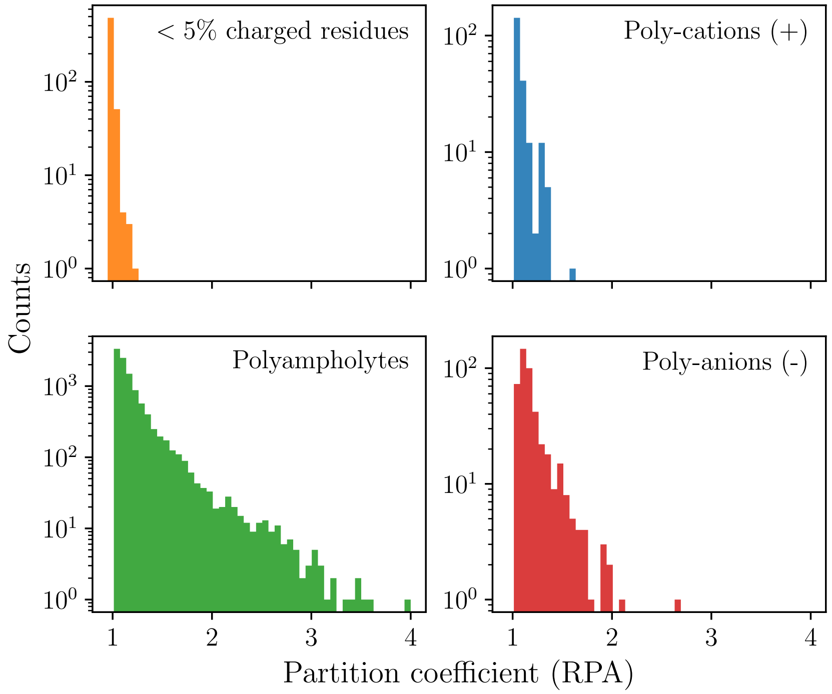


**Figure S7**: Distribution of RPA-predicted partition coefficients within the four categories of IDR sequences (as labeled in the figure) among all sequences in the Metapredict v2.4 IDRome with chain lengths $N < 4{,}100$. As stated in the text of the present Supplementary Information, using the acronyms "FCR" for fraction of charged residues and "NCCR" for net charge per charged residue, the sequences in the four categories are defined as follows: $<$ 5% charged residues: FCR $< 0.05$; Polyampholytes: FCR $> 0.2$, NCCR $< 0.2$; Poly-cations: FCR $> 0.2$, NCCR $> 0.8$; and Poly-anions: FCR $> 0.2$, NCCR $< -0.8$.

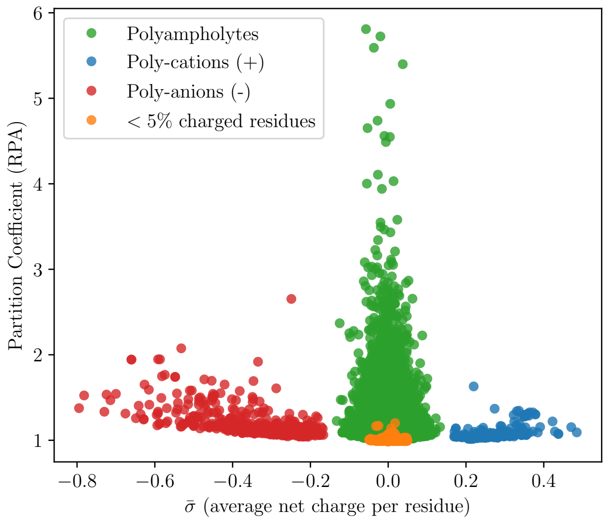


**Figure S8**: Scatter plot of RPA-predicted partition coefficient versus average net charge per residue for the four categories of IDR sequences in Fig. S7.

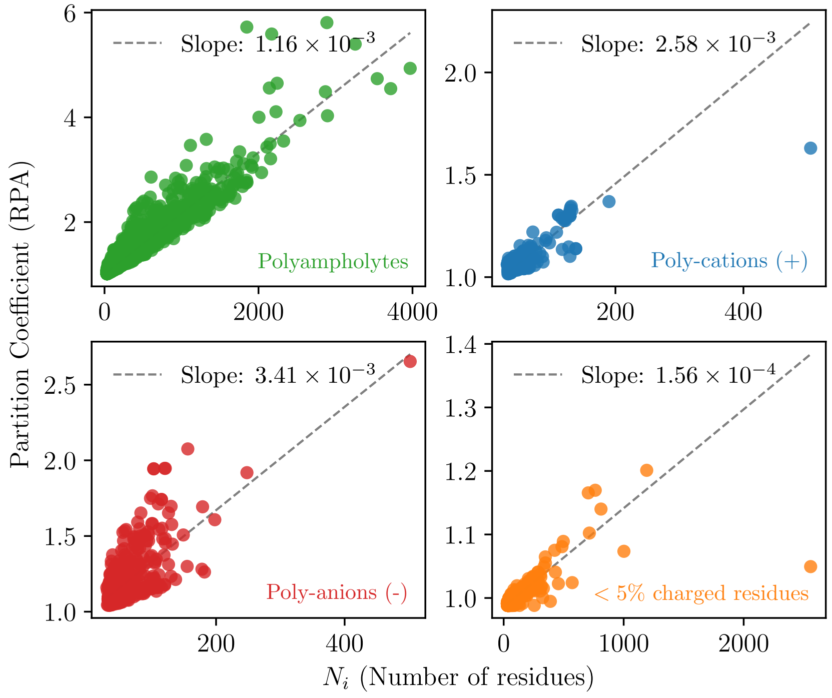


**Figure S9**: Dependence of RPA-predicted partition coefficients (PCs) on IDR chain lengths. Scatter plots show statistics of chain length dependence of predicted PCs for the four categories of IDR sequences considered in Fig. S7.

[9] Giulio Tesei and Kresten Lindorff-Larsen. Improved predictions of phase behaviour of intrinsically disordered proteins by tuning the interaction range. *Open Research Europe*, 2, 2022.
